# Inter- and intra-Subunit interactions driving the MukBEF ATPase

**DOI:** 10.1101/2022.03.08.483475

**Authors:** Soon Bahng, Rupesh Kumar, Kenneth J. Marians

## Abstract

MukBEF, an SMC-like protein complex, is the bacterial condensin. It is likely that it compacts DNA via an ATP hydrolysis-dependent, DNA loop-extrusion reaction as has been demonstrated for the yeast SMC proteins condensin and cohesin. MukB also interacts with the ParC subunit of the cellular chromosomal decatenase topoisomerase IV, an interaction that is required for proper chromosome condensation and segregation in *E. coli*, although it suppresses the MukB ATPase activity. We have investigated the MukBEF ATPase activity, identifying inter- and intra-subunit interactions by protein-protein crosslinking and site-specific mutagenesis. We show that interactions between the hinge of MukB and its neck region are essential for the ATPase activity, that the ParC subunit of Topo IV inhibits the MukB ATPase by preventing this interaction, that MukE interaction with DNA is likely essential for viability, and that interactions between MukF and the MukB neck region are necessary for ATPase activity and viability.

## Introduction

Structural Maintenance of Chromosome (SMC)^1^ proteins are essential for the management of chromosome compaction, segregation and repair. In *E. coli*, MukB, an SMC-like protein (1), acts in concert with its accessory proteins MukF, the kleisin, and MukE, a KITE (kleisin interacting winged helix tandem elements) protein (2,3) to condense the chromosome (4). Mutations in any of the genes encoding these proteins give rise to temperature sensitivity, and chromosome condensation and segregation defects (5,6). MukB also interacts with the cellular decatenase topoisomerase IV (Topo IV), mediated by amino acid residues in the hinge region of MukB and the C-terminal β propeller domain of the ParC subunit (7-9). Mutations in *mukB* that disrupt the interaction with ParC also give rise to chromosome condensation and segregation defects (10).

SMC proteins are characterized by long coiled coil arms nearly 50 nm in length that separate a head domain formed by N- and C-terminal globular regions. SMC proteins dimerize by interactions at the hinge region, which divides the coiled-coil region roughly in half (1) (Fig. 1A). The head domains of the two protomers in the dimer bind ATP between them and are responsible for the hydrolysis of ATP (11).

**Fig. 1.**
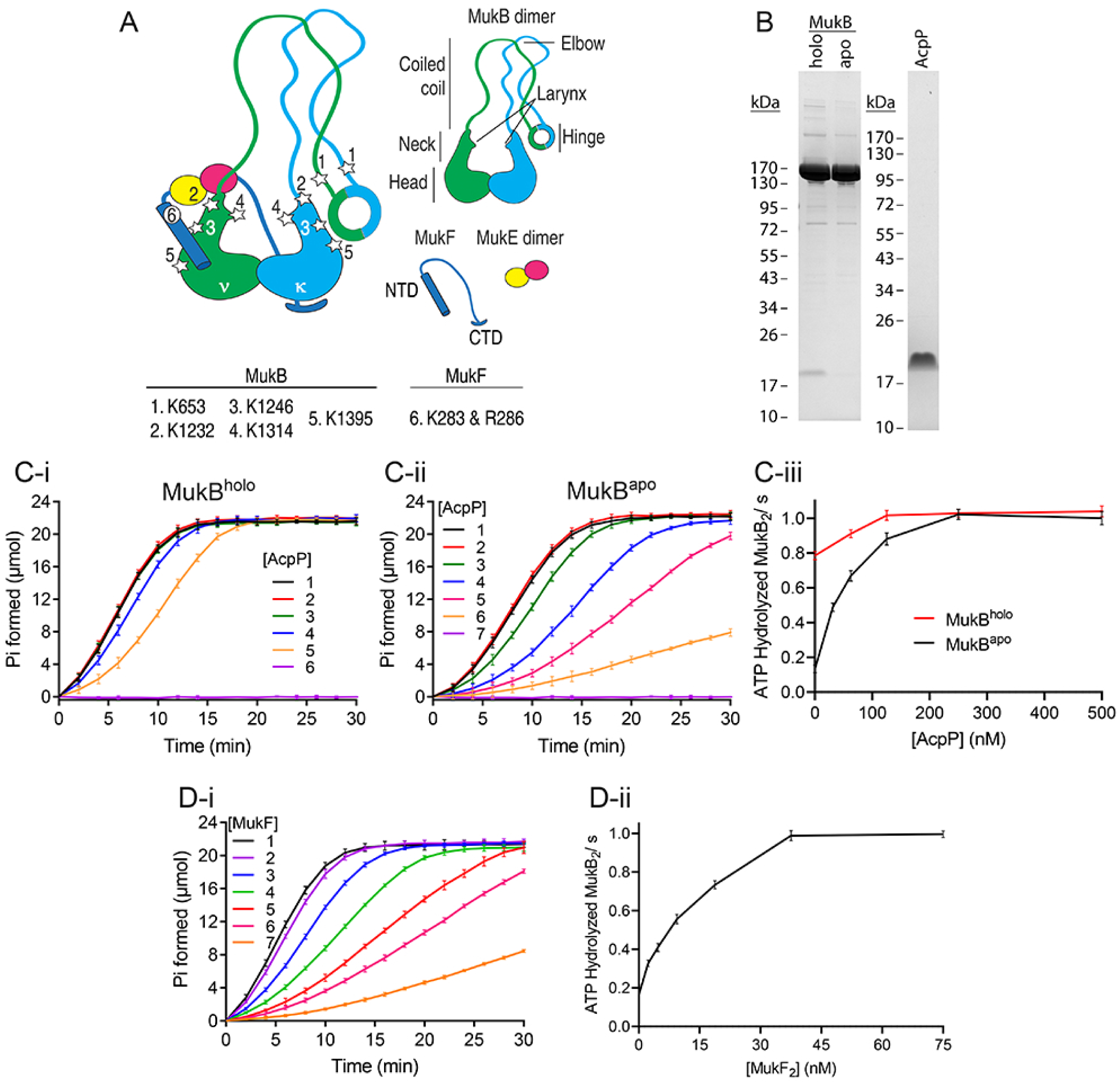
Optimization of standard conditions for MukB ATPase. A, cartoon diagram of the core MukBEF complex (after Burmann et al. (23)). AcpP has been omitted from the diagram. The two MukB dimers are denoted ν and κ. The numbered stars locate the positions of amino acid residues mutated in this study. B, SDS-PAGE analysis of MukB^holo^, MukB^apo^ (5 μg each), and AcpP (1.5 μg) through Novex 4%-20% Tris-glycine gels. C, requirement for AcpP for maximal rates of MukBF ATPase. C-i, titration of AcpP in standard ATPase reactions containing 250 nM MukB_2_^holo^ and 75 nM MukF_2_ (except reaction 6, which did not contain either MukB or MukF). In reactions 1-6 [AcpP] was 500 nM, 250 nM, 125 nM, 63 nM, none, and 500 nM, respectively. C-ii, titration of AcpP in standard ATPase reactions containing 250 nM MukB_2_^apo^ and 75 nM MukF_2_ (except reaction 7, which did not contain either MukB or MukF). In reactions 1-7 [AcpP] was 500 nM, 250 nM, 125 nM, 63 nM, 31 nM, none, and 500 nM, respectively. C-iii, plot of ATP hydrolyzed/MukB_2_/s as a function of [AcpP] from the data in panels C-I and C-ii. D-i, Titration of MukF_2_ in standard ATPase reactions containing 250 nM MukB_2_^holo^ and 250 nM AcpP. In reactions 1-7 [MukF_2_] was 75 nM, 38 nM, 19 nM, 9 nM, 5 nM, 2 nM, and none, respectively. D-ii, data from panel D-I plotted as ATP hydrolyzed/MukB_2_/s as a function of [MukF_2_].

Recently a number of SMC proteins have been shown to manipulate DNA conformation by an ATP-dependent process termed loop extrusion. Yeast condensin extrudes DNA loops in a unidirectional manner (12,13), whereas human cohesion extrudes loops bidirectionally (14,15). A “hold and feed” mechanism for loop extrusion has recently been proposed (16). Whereas a similar activity for MukBEF has not yet been reported, it has been shown that the protein complex organizes the *E. coli* chromosome into an axial core from which loops of DNA protrude (17). MukB and Topo IV are often found in the same locations on the chromosome (18-20). We have shown that when in a stoichiometric complex together, MukB and Topo IV inhibit the catalytic activities of each other: ParC inhibits the ATPase activity of MukB, whereas MukB inhibits DNA transport segment trapping and DNA cleavage by Topo IV (21), suggesting that any ATP-dependent DNA manipulation catalyzed by MukBEF is modulated by Topo IV.

Very recently a cryo-electron microscopy (cryo-EM) structure of MukBEF bound to DNA and MatP, a protein that acts to individualize the terminus region of the *E. coli* chromosome (22), revealed that the complex also contained the acyl carrier protein (AcpP) bound near the elbows of the coiled-coil regions of the two MukB protomers in a MukB dimer (23). This structure showed that the coiled-coil regions were bent so that the MukB hinge region contacted the coiled-coil region of one of the MukB protomers at the “neck,” which is just above the head domains. This bent conformation of MukB had been reported previously (24). AcpP has also very recently been shown to be required for the MukB ATPase (25).

Prior to the publication of either of the structures of MukB mentioned above, in order to understand the effect of ParC on MukB, we initiated a protein-protein crosslinking study between ParC and MukB and MukB and MukF. We report the results of this study here combined with site-specific mutagenesis of contact points within the MukB protomer, between MukB and ParC, and between MukB and MukF. We define an area within the MukB neck and head region that crosslinks with ParC and the hinge region of MukB. We show that interaction of the hinge region of MukB with its neck region is required for the MukB ATPase activity. These latter two observations indicate that ParC inhibits the MukB ATPase activity by preventing the interaction between the hinge and the neck regions. We also show that interactions between MukF and the same region of the MukB neck are required for the MukB ATPase activity and for viability and that mutation of amino acid residues of MukE that are likely to contact DNA also result in the loss of viability.

## Results

### Optimal Conditions for the MukB ATPase Activity

MukB alone hydrolyzes ATP (26), and this activity is stimulated by MukF (27,28), whereas the MukBF ATPase is inhibited by MukE (27,28). AcpP was known to interact with MukB (26,29,30), but the significance of this interaction remained obscure until Prince et al. (25) demonstrated that AcpP was required for maximal MukBF ATPase activity. This observation indicated that some previous observations of the MukB ATPase activity might have been affected by the extent to which the MukB preparation was saturated with AcpP. The cryo-EM structure of MukBEF shows the stoichiometry of AcpP to MukB to be 1:1 (23). We therefore re-investigated the optimal conditions for MukBF ATPase activity. To do so we compared the ATPase activity of a preparation of MukB that was only 5% saturated with AcpP (apo-MukB) to that of one that was nominally 100% saturated (holo-MukB) (Fig. 1).

To measure ATPase activity we used the ENZCheck Phosphate Assay Kit that detects free inorganic phosphate in the presence of 250 nM MukB_2_ and other additions to the reaction as indicated. We use least squares analysis to fit the linear portion of the phosphate accumulation curve to a straight line and calculate the rate of phosphate release. This value is then converted to ATP hydrolyzed per MukB dimer per second. AcpP was purified as described under Experimental Procedures (a gel showing the purified preparation is shown in Fig. 1B). The response of the MukBF ATPase to increasing concentrations of AcpP is shown in Fig. 1C. Even the preparation of MukB that had 1:1 stoichiometry with AcpP (based on Coomassie staining, Fig. 1B) was stimulated by additional AcpP (Fig. 1C-i). Based on this analysis, we performed all MukB ATPase assays reported herein (except those shown in Fig. 5A-ii) in the presence of 250 nM AcpP.

We had previously shown that MukF dimer saturated the MukB_2_ ATPase at a ratio of 1:4 (27). This ratio remained the same in the presence of 250 nM AcpP (Fig. 1D). We therefore selected as standard ATPase conditions 250 nM MukB_2_, 75 nM MukF_2_, and 250 nM AcpP. Assuming the majority of MukB molecules in our preparations are active (and we find this MukF saturation ratio remains the same for all MukB variants discussed in this report), this observation suggests that MukF is acting catalytically in the ATPase assay, binding to and dissociating from MukB dimers. It is known from both crystal structures of portions of MukBF (11) and from the cryo-EM structures (23) that MukF is mobile and rearranges its binding to the MukB dimer. It is therefore possible that MukF releases completely from MukB during the ATPase cycle.

### MukE Inhibition of the MukBF ATPase is Not Relieved by the Addition of DNA

The addition of MukE dimer inhibits the MukBF ATPase (27,28) (Fig. 2A). It has been reported that the addition of excess (to MukB) concentrations of small, double-stranded DNA oligomers relieved this inhibition (28). Unlike that of most SMC proteins, the MukBF ATPase is not DNA-dependent (31) and is not affected by the presence of high concentrations of a 50mer DNA duplex (Fig. 2B). We find that under the optimal conditions of MukBF ATPase, MukE inhibition was not relieved by the presence of even 10-fold more DNA oligomer than MukB (Fig. 2B). It is possible that previous observations (28) were influenced by sub-stoichiometric concentrations of AcpP in the MukB preparations used.

**Fig. 2.**
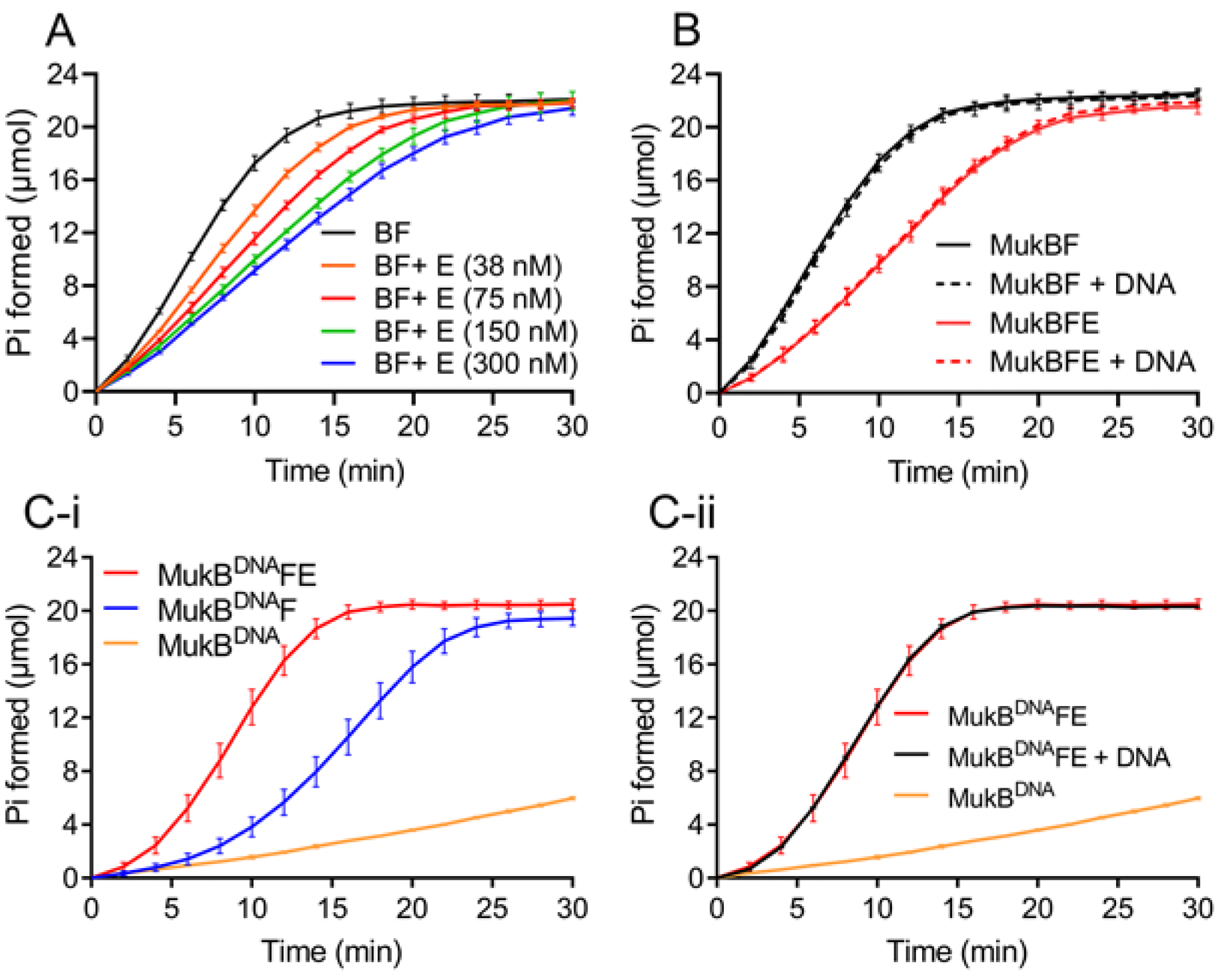
DNA does not relieve the inhibition of the MukBF ATPase by MukE. A, Standard ATPase reactions contained the indicated concentrations of MukE_4_. B, DNA does not relieve the inhibition of the MukBF ATPase by MukE. Standard ATPase reactions contained 300 nM MukE_4_ and 2.5 μM of a 50 bp double-stranded oligonucleotide as indicated. C-i, The ATPase activity of MukB^DNA^F is stimulated by MukE. Standard ATPase reactions contained MukB^DNA^, MukB^DNA^F, or MukB^DNA^F with 300 nM MukE_4_ as indicated. C-ii, DNA has no effect on the MukB^DNA^FE ATPase. Standard ATPase reactions as indicated. DNA, when present, was as in panel B.

The cryo-EM structure of MukBEF shows, however, that MukE forms part of the clamp that binds DNA to the MukB heads (23). To address the effect of DNA binding to MukB on MukE modulation of the MukBF ATPase activity, we used a MukB variant we had previously developed (27), MukB R187E R189E (MukB^DNA^), that binds DNA with one-quarter the affinity of the wild-type protein and manifests a reduced ATPase activity compared to the wild type (compare Figs. 2A and 2C-i). Remarkably, MukE stimulated the ATPase of this protein, rather than inhibited (Fig. 2C-i). Added DNA had no effect (Fig. 2C-ii). The decreased ATPase activity of MukB^DNA^ suggests that the mutated MukB residues perturb necessary inter-subunit interactions and that this perturbation could be overcome to some extent by the presence of MukE, likely by a stabilizing effect on the MukBEF complex. Therefore, to address the role of DNA in the ATPase cycle, we turned our attention to MukE, which makes a significant number of interactions with the DNA bound to the head in the presence of ATP (23).

In the structure of the core MukBEF complex (MukBEF, AcpP), MukE resides outside the MukB lumen and, as the MukBEF holo-complex (MukBEF, MatP, DNA, and ATP) is assembled with DNA and ATP, it moves closer to the bound DNA and forces open the neck of νMukB. This DNA binding concurrent with the opening of the neck by MukE, points to its important role in the ATPase reaction. We made charge-reversal mutations in six positively charged residues (R78E, R85E, K150E, R156E, R163E and R164E; MukE^6X^), based on their proximity to the DNA in the structure.

MukE^6X^ was significantly less inhibitory to the MukBF ATPase than wild-type MukE in either the absence or presence of DNA (Figs. 3A and B). Examination of the crystal structure of fragments of MukBEF (11) suggests that in the absence of DNA, MukE could enter even further into the MukB lumen than it does in the cryo-EM structure (23) to sit at the top of the MukB heads with MukE residues R78 and R85 proximal to the positively charged DNA-binding region in the MukB head domain. Charge reversal substitutions at these residues may therefore stabilize MukE in this position, favoring an ATP-engaged conformation of the MukB heads to stimulate ATP hydrolysis.

**Fig. 3.**
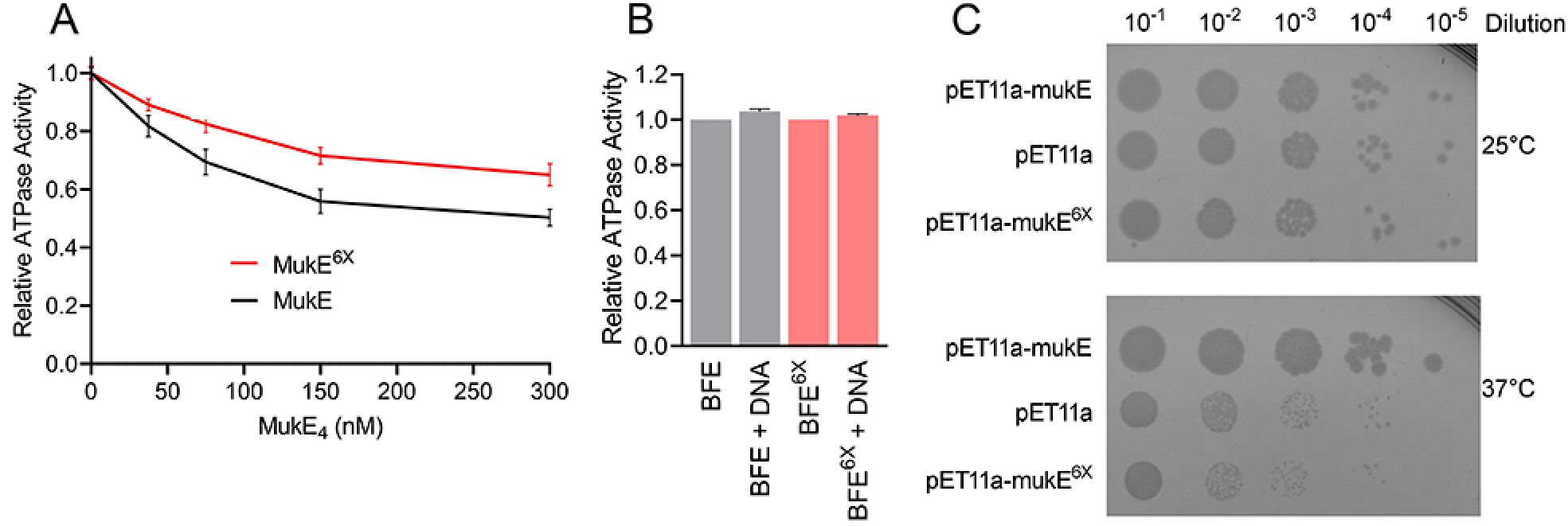
The MukE^6X^ variant is less inhibitory than the wild type to the MukBF ATPase. A, the graph shows the relative maximal ATPase activity determined in standard ATPase reactions containing either wild-type or MukE^6X^ at the indicated concentrations. B, DNA has no effect on the MukBFE^6X^ ATPase activity. Standard ATPase reactions contained either wild-type or MukB^6X^ and DNA (as in Fig. 2B) as indicated. C, the *mukE*^*6X*^ allele does not complement a temperature-sensitive a *mukE::kan* strain. AZ5450 cells transformed with the indicated plasmids were grown to mid-log phase at 25 °C, serially diluted, plated on LB plates containing ampicillin, and grown at 25 °C and 37 °C.

Because the DNA is clamped between MukB, MukE, and MukF, any alterations to the DNA binding residue are likely to affect the dynamics of the MukBEF complex in the absence of DNA. Although the ATPase activity is not affected by DNA binding in the assays reported here, the energy generated from ATP hydrolysis is believed to fuel translocation of the complex on DNA. To address the connection between the two activities in overall function of the MukBEF complex, we carried out complementation assays with plasmids carrying *mukE* and *mukE*^*6X*^ alleles using a temperature-sensitive strain where *mukE* was disrupted (Fig. 3C). Whereas wild-type *mukE* complemented well at the non-permissive temperature, *mukE*^*6X*^ failed to do so, underscoring the importance of these DNA binding residues in vivo. Expression levels of the *mukE* variants from the plasmids was equivalent (Fig. S1).

### Protein-Protein Crosslinking Reveals Regions of the MukB Head and Neck that Interact with the MukB Hinge Region, MukF, and ParC

In order to develop insight into the interactions between MukBEF subunits and the mechanism of inhibition of the MukBF ATPase by the ParC subunit of Topo IV (21), we performed disuccinimldyl glutarate-catalyzed protein-protein crosslinking experiments with MukB alone, MukB and ParC, and MukB and MukF, followed by trypsin digestion and analysis of the cross-linked peptides by mass spectrometry. The crosslinks found between ParC and MukB, MukF and MukB, and selected crosslinks for MukB alone are displayed in Fig. 4. Complete crosslinking results are included as Tables S1-S3.

**Fig. 4.**
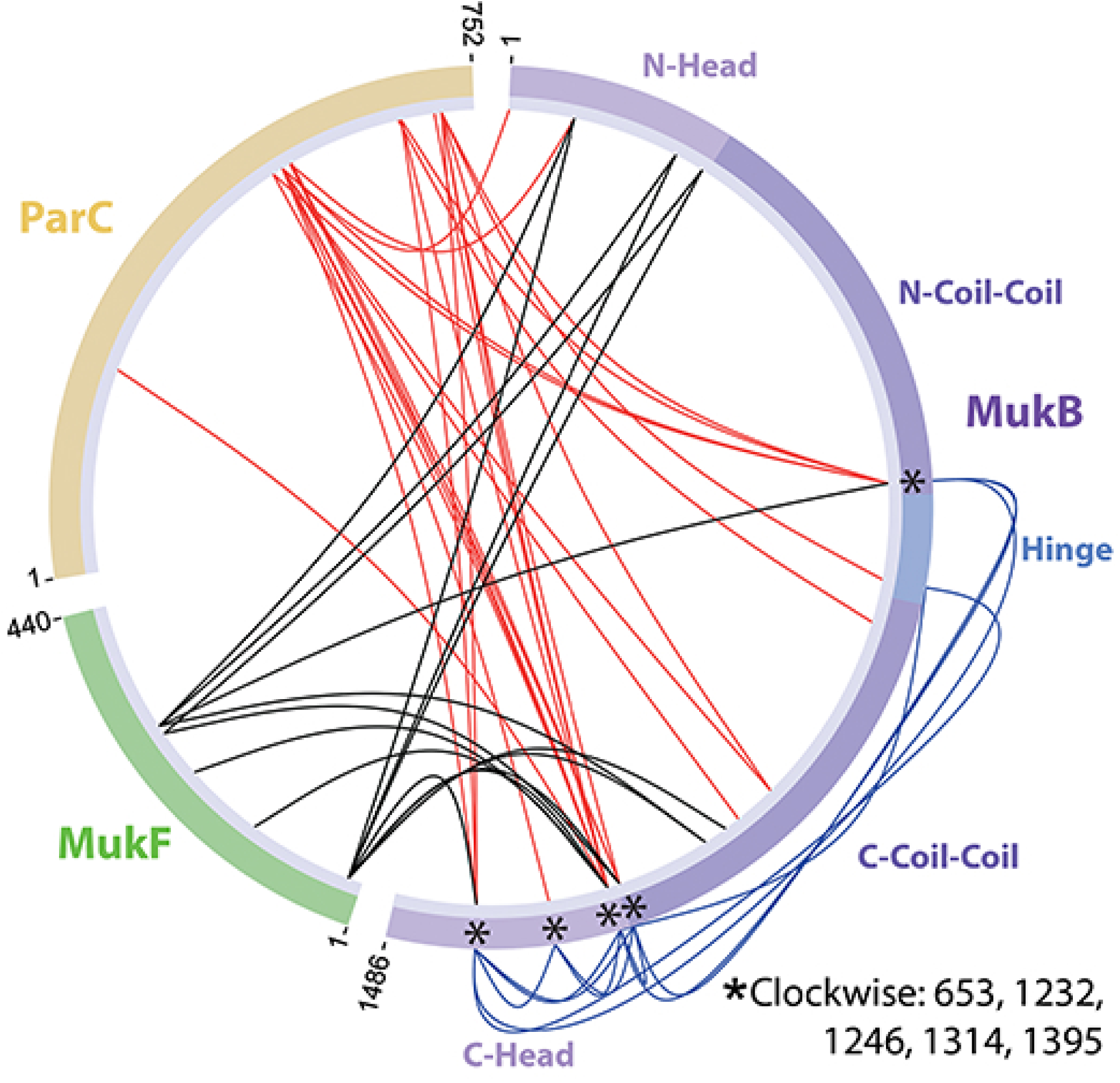
Protein-protein crosslinking results between MukB-MukB, MukB-MukF, and MukB-ParC. Protein-protein crosslinking was performed and analyzed as described under Experimental Procedures and are presented in circular format using XiView software (47). Asterisks represent amino acid residues of MukB common to all three crosslinking analyses. Complete crosslinking results are provided in Tables S1-S3.

The long MukB protomer has globular domains at its N- and C-terminal ends that form half of the ATPase head domain in a MukB dimer. The coiled-coil domains are divided in half by another globular domain, the hinge, which dimerizes two MukB monomers to form the MukB dimer. Both MukB monomers in the MukB dimer fold over at a region called the elbow, resulting in the hinge domain becoming juxtaposed with the neck region, constituted of coiled-coils just above the head domain, of one of the MukB monomers, κ-MukB. The C-terminus of one MukF monomer in the MukF dimer binds to the cap region of κ-MukB. A long linker domain of MukF then snakes across the head domain of the ν-MukB monomer with its middle domain (MD) making contacts with the neck of the κ-MukB monomer, forming a clamp for DNA bound in the MukB DNA-binding domain in the joined MukB head domains. The N-terminus of MukF is bound to the head domain near the DNA binding site in the dimeric structure. The N-terminal domains from two dimeric MukB complexes interact to assemble a dimer of dimers (11,32). Two MukE dimers are bound across the winged helix domain of MukF forming part of the DNA clamp and making contacts with the ν-MukB neck domain (Fig. 1A) (11,23,24).

Our crosslinking studies defined three major regions on the outer surface of each MukB monomer marked by residues K1232 and K1246 on the neck, and K1395 on the head domain that interacted with ParC, MukF, and the MukB hinge (Fig. 4). These interactions are not directly with the catalytic ATPase center and are therefore likely to play a regulatory role, as shown below. Several MukB amino acid residues in the neck crosslinked with the hinge region and MukF, reflecting the asymmetric disposition of MukF across the two protomers of the MukB dimer and the manner in which the coiled-coil regions are folded over. Surprisingly, ParC also crosslinked with residues in both the hinge, neck, and head regions, leading us to investigate the role of the MukB hinge region in the MukB ATPase cycle.

### Interaction Between the MukB Hinge and Neck Regions Drives the MukB ATPase Cycle and ParC Binding Disrupts this Interaction

We and others had demonstrated previously that the fifth blade of the ParC C-terminal β propeller domain interacts with a region of MukB where the coiled-coils extend from the hinge (7-9). Neither the variant ParC R705E R729A nor MukB D697K D745K E753K interacts with either wild-type MukB or ParC, respectively, in pull down assays. Replacement of *mukB* with the *mukBD697KD745KE753K* allele results in severe nucleoid decompaction and chromosome segregation defects in vivo (10).

ParC inhibits the MukBF ATPase (21), implying, given the folded conformation of MukB (23,24) and our crosslinking results, that ParC might interpose itself between the MukB hinge and neck regions and that this might disrupt the ATPase cycle, further suggesting that isolated MukB hinge might stimulate the MukBF ATPase *in trans*. This proved to be the case.

The addition of purified MukB hinge [MukB amino acid residues 645-804 (33)] had little effect on the ATPase activity of MukB under our optimal conditions in the presence of 250 nM AcpP (Fig. 5A-i). However, the MukB hinge had a significant stimulatory effect in the absence of AcpP on the activity of apo-MukB compared to holo-MukB that was in 1:1 complex with AcpP (Figs 5A-ii and A-iii). We had previously shown that addition of purified MukB hinge to full-length MukB bound to DNA caused dissociation of the latter protein from the DNA (33). In order to rule out the possibility that our MukB hinge preparation was contaminated with AcpP, we analyzed both trypsin and chymotrypsin digests in solution of large amounts of the MukB hinge preparation by mass spectrometry. Whereas other trace contaminants were detected, no AcpP-derived peptides were detected at all (Table S4). We conclude that the addition of the MukB hinge *in trans* can stimulate the ATPase activity of the intact protein, suggesting that contact between the hinge and the neck regions of MukB forms part of the ATPase cycle.

**Fig. 5.**
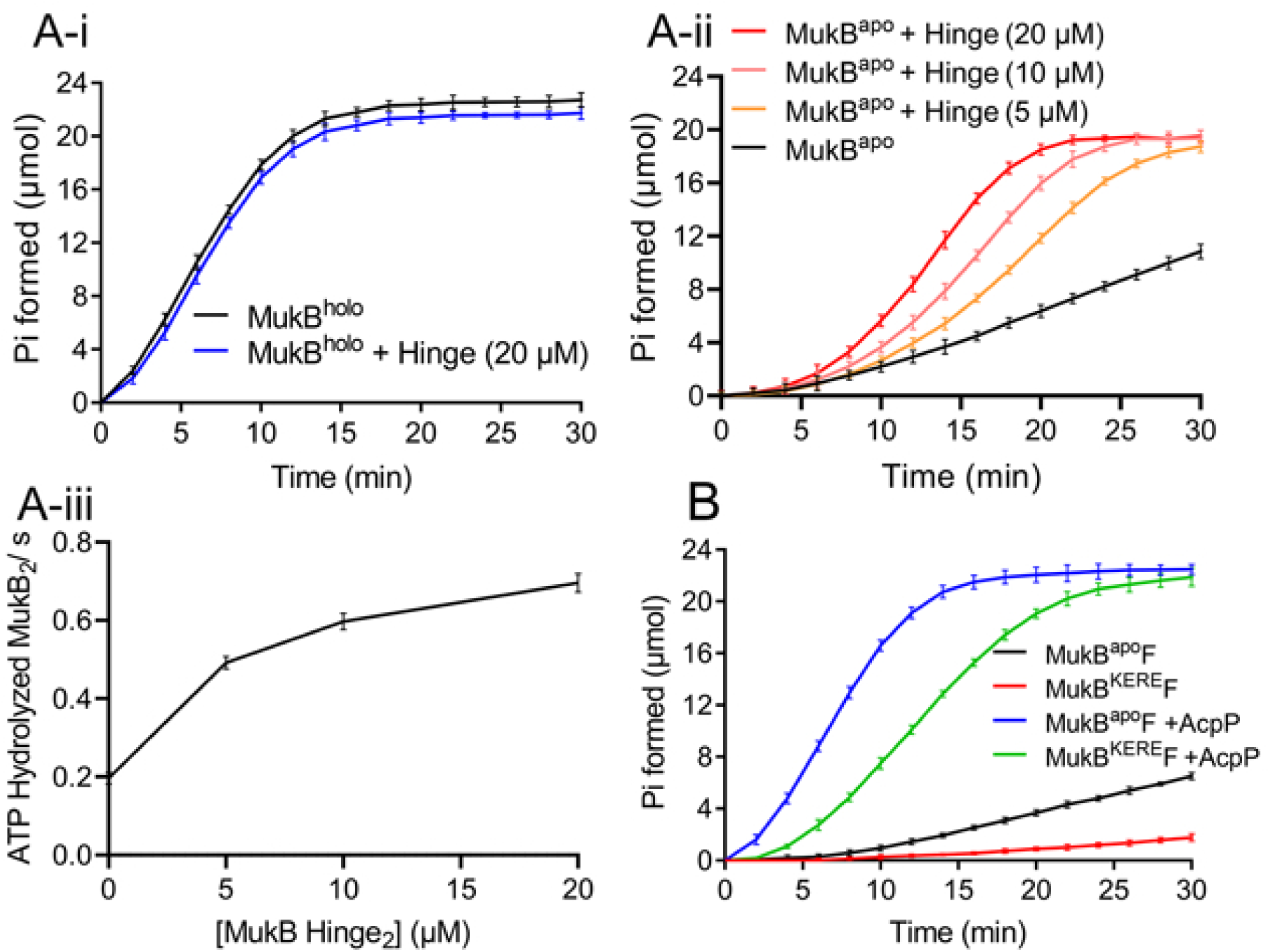
The MukB hinge fragment stimulates the ATPase activity of MukB^apo^ *in trans*. A-i, purified MukB hinge fragment does not stimulate the ATPase activity of MukB^holo^. Standard ATPase reactions with MukB^holo^F either did or did not contain 20 μM purified MukB hinge_2_ fragment as indicated. A-ii, purified MukB hinge fragment does stimulate the ATPase activity of MukB^apo^. Standard ATPase reactions with MukB^apo^F either contained no MukB hinge_2_ fragment or did contain the indicated concentrations of MukB hinge_2_ fragment. A-iii, the data in panel A-ii converted to ATP hydrolyzed/MukB_2_/s and plotted as a function of the concentration of MukB hinge_2_ fragment. B, the full-length MukB hinge mutant, MukB^KERE^, has significantly reduced ATPase activity compared to wild-type MukB. Standard ATPase reactions contained either wild-type MukB^apo^F or MukB^KERE^F in the presence or absence of AcpP as indicated.

We noted that our crosslinking results showed interaction between amino acid residues K761 and K1395 in the MukB hinge and head region, respectively. We had previously shown that strains carrying the *mukBK761ER765E* allele were elongated and had nucleoids of increased size compared to wild type (33). These mutations are in the hinge region and include the residue that crosslinks to the neck region. The MukB variant K761E R765E (MukB^KERE^) has significantly reduced ATPase activity compared to wild type even in the presence of saturating concentrations of AcpP (Fig. 5B), further emphasizing the importance of the MukB hinge-neck/head contact in the ATPase cycle.

### *A MukF Variant Defective in Stimulation of MukB ATPase and* mukF *Complementation*

Our MukF-MukB crosslinking (Fig. 4) showed crosslinking between MukF residues K283 and K293 with MukB neck residue K1232. Based on these results, we mapped these residues on available MukF and MukB crystal structures (11) before the publication of the MukBEF cryo-EM structure (23). Mapping of K283 on the MukF structure, followed by the generation of local protein contact potential using PyMOL (v. 2.0, Schrödinger, LLC), showed it to be part of a basic patch. On MukB, the residue corresponding to K1232 was not present in the crystal structure, which started at I1235. This residue was also located in a basic patch, however, this basic patch was flanked by two acidic patches on either side. We therefore assumed an electrostatic interaction between the basic and acidic patches on MukF and MukB, respectively. Such an interaction could place the two crosslinking residues (MukB K1232 and MukF K283) near each other. To address this interaction and its importance in the function of MukBEF complex, we replaced K283, along with another residue, R286, present in the same basic patch, with glutamates, generating MukF^2X^. MukF^2X^ was 40%-50% less active than wild-type MukF in stimulating the MukB ATPase (Fig. 6A). Interestingly, when provided *in trans* on a plasmid, *mukFK283ER286E* failed to complement the temperature sensitivity of a *mukF* disruption strain, whereas the wild-type allele complemented completely (Fig. 6B). Both proteins were expressed to the same level (Fig. S1)

**Fig. 6.**
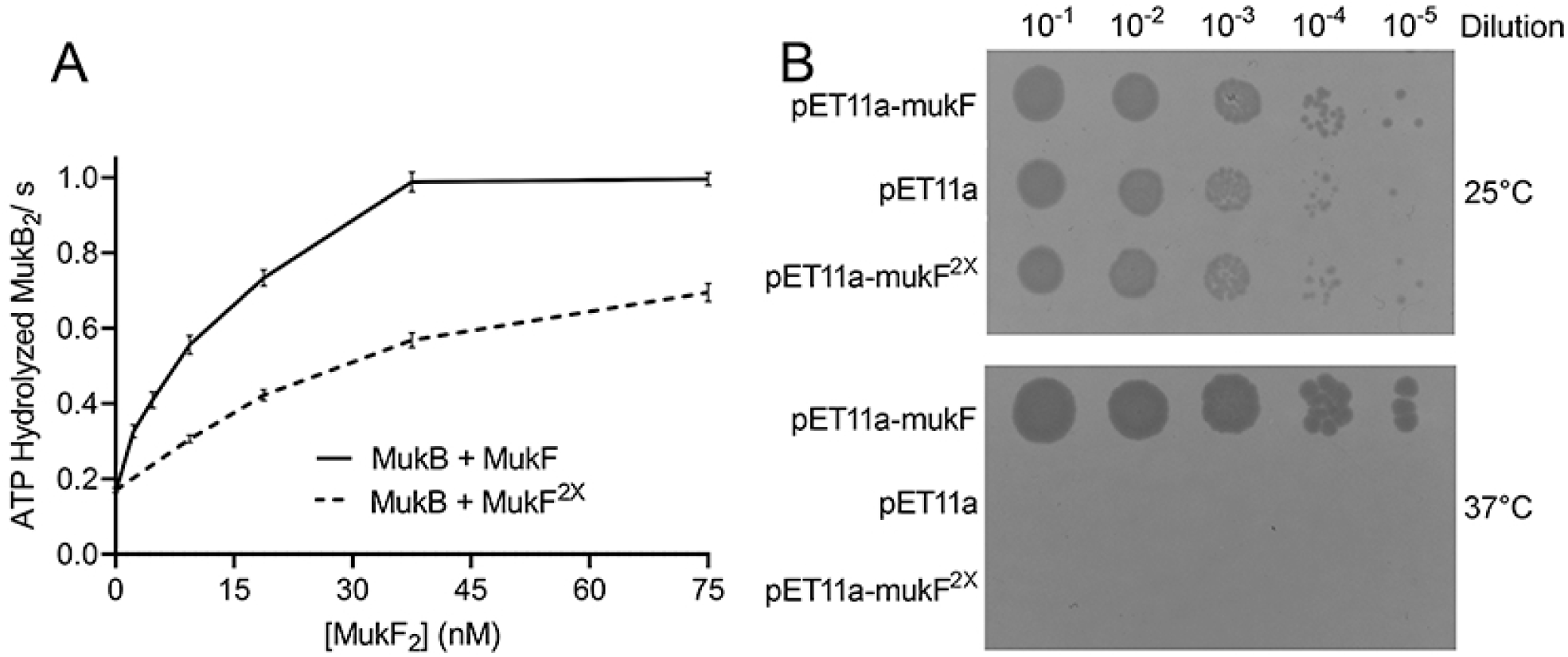
The MukF^2X^ variant is less active than wild type in stimulating MukB ATPase activity. A, maximal rates of ATPase activity from standard ATPase reactions containing either no wild-type MukF_2_ or MukF_2_^2X^ or 2 nM, 4.7 nM, 9.4 nM, 18.9 nM, 37.5 nM, and 75 nM wild-type MukF_2_ or MukF_2_^2X^ plotted as a function of [MukF_2_]. B, the *mukF*^*2X*^ allele does not complement the temperature sensitivity of a *mukF::kan* strain. AZ5381 cells transformed with the indicated plasmids were grown to mid-log phase at 25 °C, serially diluted, plated on LB plates containing ampicillin, and grown at 25 °C and 37 °C.

### MukB Variants Mutated at Head and Neck Residues Are Deficient in MukF Stimulation and MukE Inhibition of the ATPase Activity

We engineered alanine replacement and charge reversal variants of MukB in the amino acid residues we found in the head and neck regions that crosslinked to the hinge region, ParC, and MukF. Under our optimal conditions for ATPase activity, all ten MukB variants had reduced maximal rates of ATP hydrolysis compared to the wild-type (Table 1).

**TABLE 1:** Maximal ATPase rates of wild type and mutated MukBvariants

| MukB Variants | ATP hydrolyzed MukB<sub>2</sub> / s |
| --- | --- |
| wildtype | 0.99 + 0.07 |
| K761ER765E | 0.62 + 0.01 |
| R187ER189E | 0.56 + 0.01 |
| K653A | 0.69 + 0.01 |
| K653E | 0.64 + 0.02 |
| K1232A | 0.79 + 0.01 |
| K1232E | 0.72 ± 0.01 |
| K1246A | 0.67 + 0.07 |
| K1246E | 0.73 + 0.03 |
| K1314A | 0.71 + 0.02 |
| K1314E | 0.74 + 0.01 |
| K1395A | 0.68 + 0.02 |
| K1395E | 0.67 + 0.02 |

We examined the ATPase activity of these variants with respect to stimulation by MukF and MukF^2X^, inhibition by ParC, and inhibition by MukE. Crosslinking showed that all five of the MukB amino acid residues mutated interacted with ParC (Fig. 4), we therefore examined the inhibition of the MukBF ATPase by ParC for all of the ten variants (Fig. 7). There was no significant difference in ParC inhibition observed. This result is not surprising given that the primary site of interaction between MukB and ParC is the hinge domain, allowing ParC to prevent MukB hinge-neck and hinge-head interactions despite mutations at these sites. Therefore, the crosslinking results presumably reflect the disposition of ParC once it has interposed itself between the head/neck and hinge regions of MukB

**Fig. 7.**
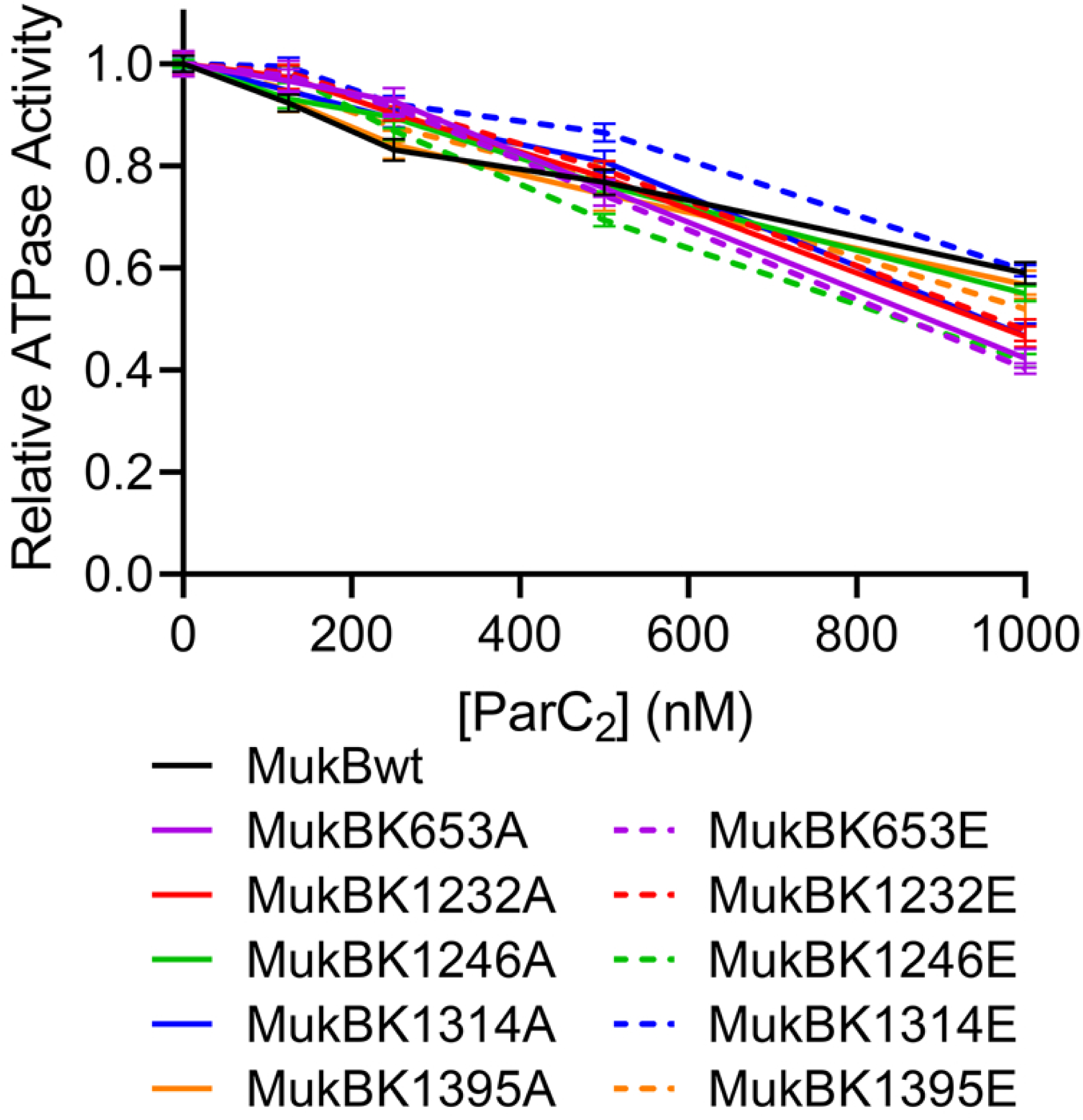
The ATPase activities of MukB head and neck variants are equally responsive to inhibition by ParC as wild-type MukB. The ATPase activities of MukB wild type and the indicated MukB variants were determined in the presence of either no or 125 nM, 250 nM, 500 nM and 1 μM ParC. Results are presented for each MukB protein as activity relative to the rate of ATP hydrolysis in the absence of ParC.

Four of five of these amino acid residues interact with MukF, presumably reflected in the decrease in maximal ATPase rates shown in Table 1. Given these established interactions we determined whether any of the variants were differentially sensitive to stimulation by MukF^2X^ compared to wild-type MukF. To do so we plotted the difference between MukF wild type and MukF^2X^ in ATP hydrolysis elicited at equivalent concentrations of the proteins (Fig. 8). Most of the variants did not show any difference in sensitivity to MukF^2X^ compared to wild-type MukB. However, amino acid substitutions at K1246 of MukB showed clear and remarkably opposite effects (Fig. 8C): MukB K1246E was less responsive to MukF^2X^ stimulation than the wild type, whereas MukB E1246A was more responsive to MukF^2X^ stimulation than the wild type.

**Fig. 8.**
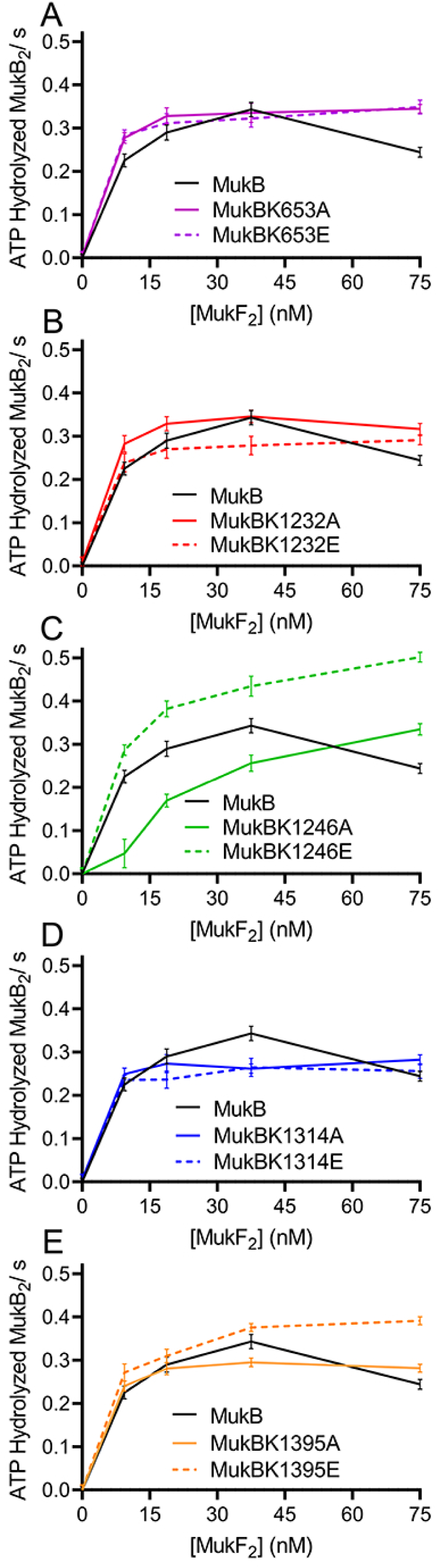
The MukB variants K1246A and K1246E respond differently to stimulation by MukF^2X^ compared to wild-type MukF. The MukF- and MukF^2X^-stimulated ATPase activities of wild-type MukB and all ten MukB head and neck variants were compared by plotting the difference in the rate of ATP hydrolysis elicited between wild-type MukF and MukF^2X^ either in the absence of MukF proteins or in the presence of 9 nM, 19 nM, 38 nM, and 75 nM MukF_2_ proteins. A, MukB K653A and MukB K653E. B, MukB K1232A and MukB K1232E. C, MukB K1246A and MukB 1246E. D, MukB 1314A and MukB 1314E. E, MukB 1395A and MukB 1395E. The curve for MukB wild type is reproduced on each graph for comparison.

The importance of neck residue K1246 also emerged when the extent of inhibition by MukE of the ATPase activity of the variants was examined (Fig. 9). Both MukB K1246A and K1246E were more sensitive to MukE inhibition than the wild type (Fig. 9C), as were the MukB K653A and K653E (Fig. 9A), and MukB K1395A and K1395E (Fig. 9E) pairs. The K653 residue is in the hinge region of MukB whereas, as noted above, the K1395 residue crosslinks with the K716 residue in the hinge region. In the Burmann structures (23), MukE is closer to νMukB, whereas MukB residues K1395 and K653 are likely involved in the interaction between the head and neck domains of κMukB on the opposite side. Therefore, the differential responses to MukE inhibition described above indicate an overall connection between the interactions of the MukB hinge and MukF and of MukE with the MukB head and neck residues on either side.

**Fig. 9.**
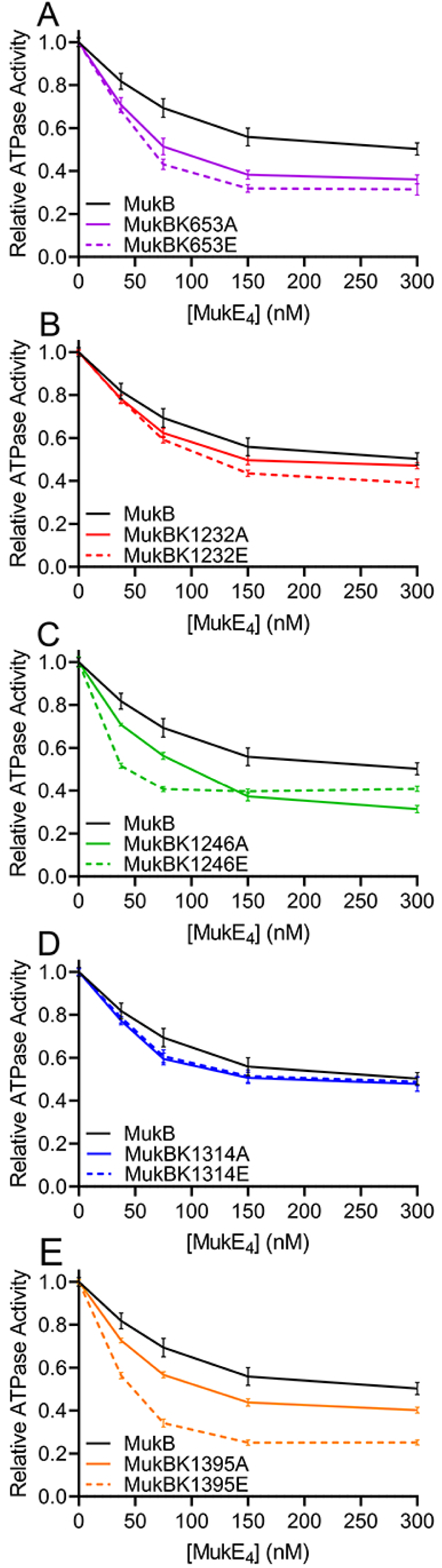
The effect of MukE on the ATPase activities of the ten MukB head and neck variant proteins. The maximal rates of ATPase activities for the ten variant proteins in either the absence or presence of 38 nM, 75 nM, 150 nM, and 300 nM MukE were determined and are presented for each MukB protein as activity relative to the rate of ATP hydrolysis in the absence of MukE. A, MukB K653A and MukB K653E. B, MukB K1232A and MukB K1232E. C, MukB K1246A and MukB 1246E. D, MukB 1314A and MukB 1314E. E, MukB 1395A and MukB 1395E. The curve for MukB wild type is reproduced on each graph for comparison.

Of the ten variants discussed above, only the *mukBK1314E* allele showed a significantly reduced extent of complementation of the temperature sensitivity of a *mukB* deletion strain (Fig. 10), despite being expressed to similar levels as the wild-type protein (Fig. S1) We had previously shown the importance for interaction with ParC of residues in the MukB hinge domain. MukB K1314 also crosslinks with ParC. The reduced complementation of the mutant allele therefore reveals a more elaborate interaction between ParC and MukB that includes not only the MukB hinge, but also extends to the MukB larynx that sits above the DNA-binding region in the MukB head domains. We consider the implications of these data in the Discussion.

**Fig. 10.**
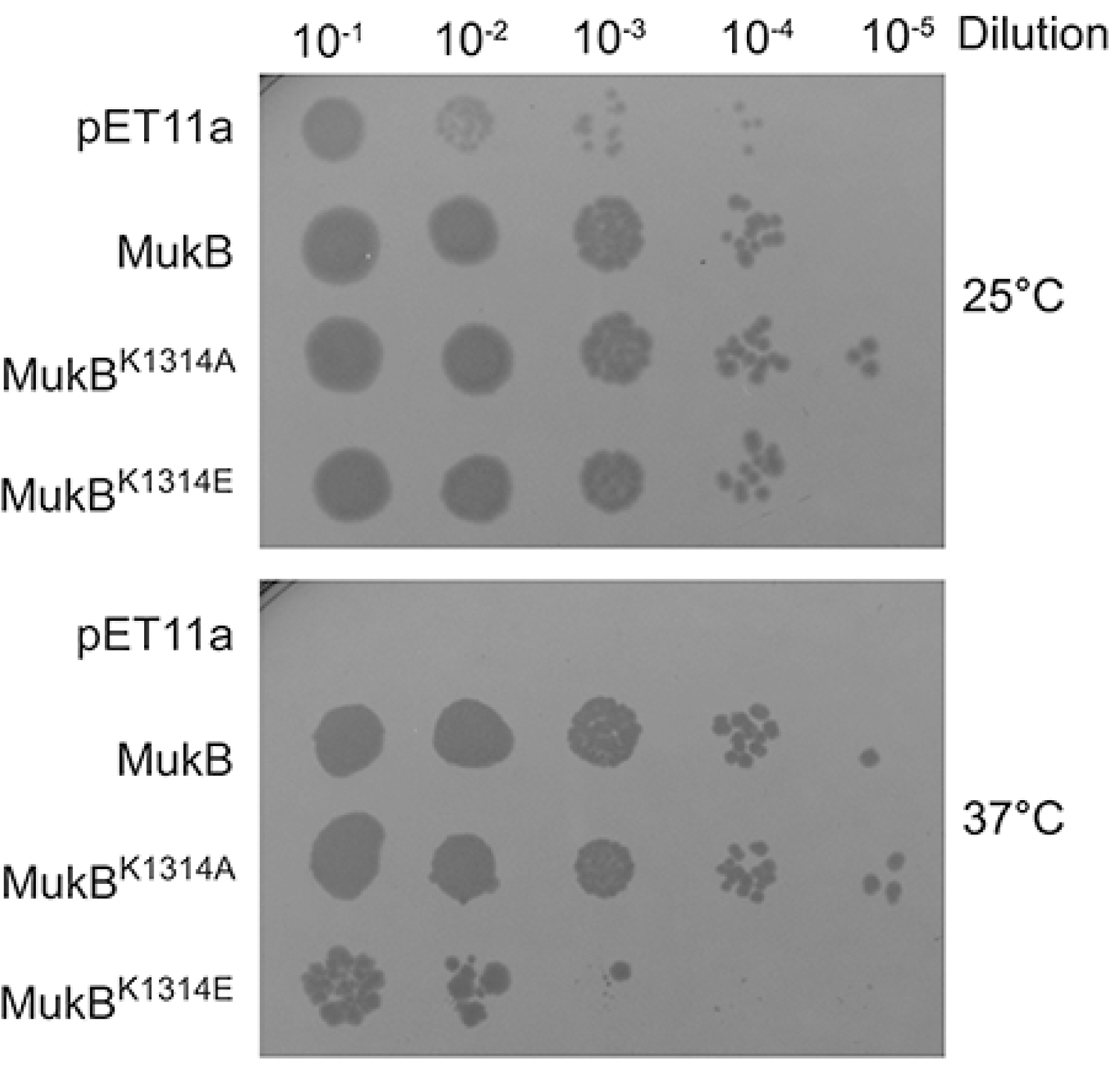
The *mukB1314A* allele is less active than the wild-type allele in suppressing the temperature sensitivity of a *ΔmukB* strain. BW30270*ΔmukB::kan* cells transformed with the indicated plasmids were grown to mid-log phase at 25 °C, serially diluted, plated on LB plates containing ampicillin, and grown at 25 °C and 37 °C.

## Discussion

SMC complexes have been referred to commonly as ring-shaped multiprotein complexes (1). SMC dimers themselves adopt a ‘V’ shape (34) that is formed into a ring by association of the kleisin and kleisin interacting proteins (3). The ring-shaped organization of these complexes suggested that DNA could become trapped topologically within the ring (35), a property that has been demonstrated experimentally for nearly every SMC complex (36). The presence of an ATP binding cassette (ABC) ATPase motif in the head domains of the SMC proteins and the architectural similarity of these domains with those in the cytoskeletal motor proteins myosin, kinesin, and dynein suggested a motor activity for the SMC complexes enabling them to translocate actively on DNA. However, how an open ring-shaped structure might translocate on DNA was difficult to envision. Indeed, a study aimed at understanding the translocation of cohesin complexes suggested that the ring of a cohesin complex may not exist in an entirely open conformation (37). Several other reports examining the topology of the SMC complexes also pointed to an alternate organization. The role of the hinge domain in DNA loading and unloading reactions of the *B. subtilis* SMC complex indicated a bending of the coiled coil that would position the hinge near the head domain (38). A recent report demonstrated that condensins, cohesins, and MukBEF all adopted a folded conformation (24). Instead of a ring shape, the coiled coil regions of the SMC dimers aligned from hinge to the head leaving only a small opening near the head. Furthermore, the coiled coil region bent to position the hinge near the head domain. It has been suggested that such an organization prevails in vivo and is likely the active form of the complex. However, the importance of such a conformation in the function of these SMC complexes, specifically their ATPase activity, has remained enigmatic.

In this study, we have uncovered intra- and inter-molecular interactions between the subunits of the MukBEF complex that determine its ATP hydrolysis activity. We show, although the head domain of MukB, like other SMC proteins, contains the residues for binding and hydrolysis of ATP at their dimerization interface, the process of hydrolysis involves dynamic interactions between MukF and the MukB hinge domain with the outer surface of the head domain of MukB. We also present a biochemical basis for the inhibition of this activity by the MukB interacting protein ParC, a subunit of topoisomerase IV. ParC, by interacting with both the MukB hinge and head domains, inhibits the ATPase activity likely by blocking interaction between the MukB hinge and head domains.

### A Push by the Hinge Domain at the Head and Neck of MukB is Required for ATPase Activity

Our crosslinking results with MukB alone suggest that it is organized in a folded conformation not requiring interaction with either MukE or MukF. Previous reports had shown that the complete MukBEF complex was in the folded conformation (23,24). Although the fraction of MukB existing in a folded conformation in solution at any given time is unknown, given that folding occurs at a specific region, the elbow, on the coiled coil, we assume that the protein may be able to readily switch between a folded and more extended conformation. Three residues, K1232, K1246 (neck) and K1395 on the head that crosslink with the hinge are located on the outer surface of the protein such that the hinge could interact with them when the coiled-coils are folded without disrupting the association of head domains (Figs. 1A and 11A). Contrary to the minor perturbation in the interaction that one would expect from mutating these residues, a significant reduction was observed in the ATPase activity for each of these mutants, indicating a critical role for these interactions. The importance of the head-hinge interaction was further underscored by the ability of the isolated hinge domain to stimulate the ATPase activity *in trans* of a preparation of MukB depleted of AcpP. The role of the hinge-head interaction in the MukB ATPase activity is further reinforced by the manner in which ParC inhibits the ATPase activity.

**Fig. 11.**
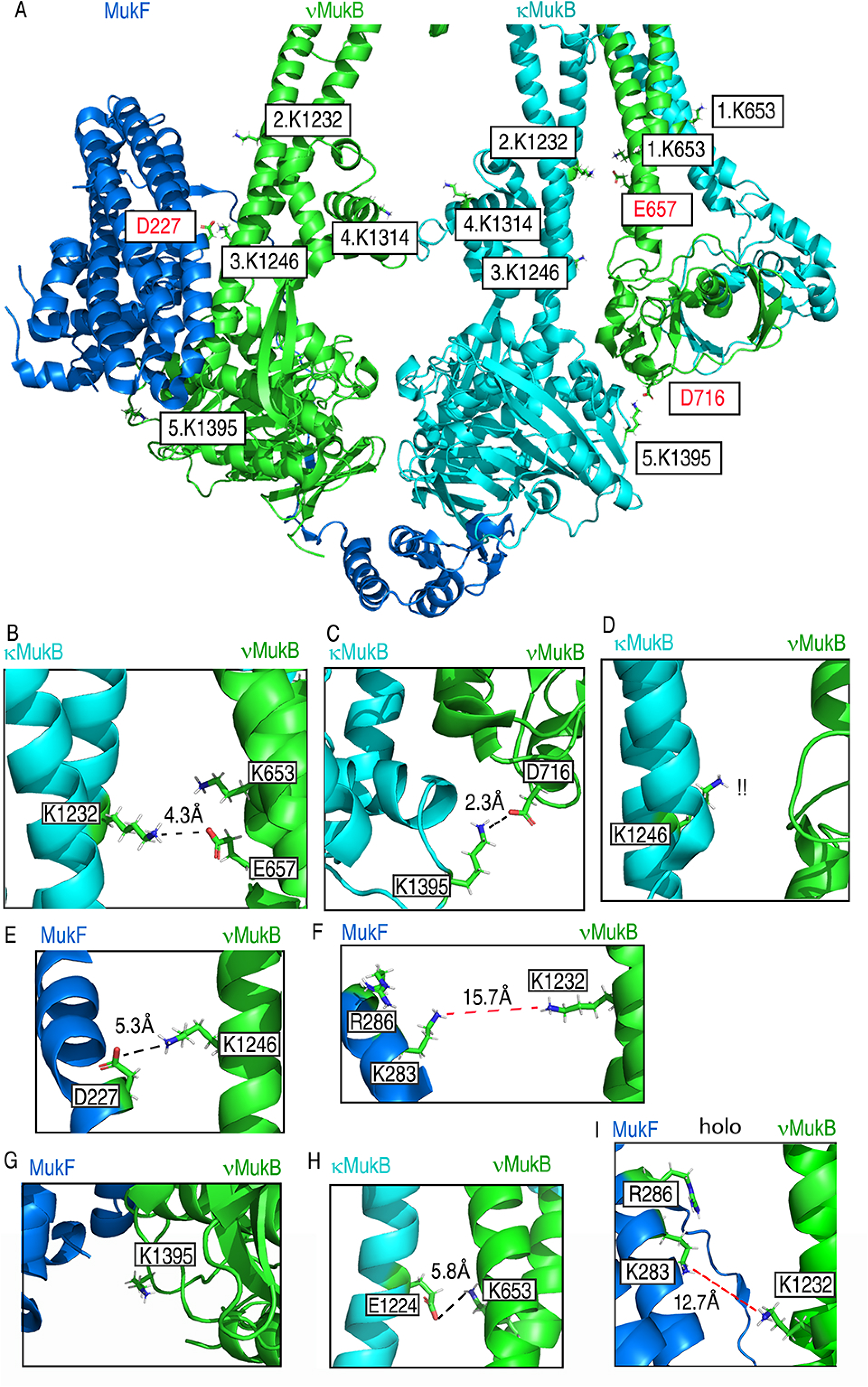
Mapping of MukB head and neck mutations to the MukBEF structures of Burmann et al. (23). A, overall view of the head, neck and larynx regions of the MukBF dimer in the core MukBEF cryo-EM structure. Amino acid residues discussed in the text are indicated. κMukB, cyan; νMukB, green; MukF, blue. B-I, specific potential interactions as discussed in the text. Panels B-H are extracted from the core MukBEF structure, whereas panel I is extracted from the holo-MukBEF structure.

### A Biochemical Basis for ParC-mediated Inhibition of the MukB ATPase Activity

Mutation of either residues D697, E745, and E753 in the MukB hinge or residues R705 and R729 in the ParC C terminal domain disrupt the interaction between MukB and ParC (7,8). MukB and ParC variants carrying mutations in these mutations show no detectable interaction between the two proteins in pulldown assays, suggesting the MukB hinge as the only interaction site for ParC. However, here, based on protein crosslinking, we report a large interface between ParC and MukB spanning the coiled coil, neck, and head domains, in addition to the sites reported previously on the hinge domain. Moreover, the interaction site of ParC on the MukB head and neck overlaps with those identified for the MukB hinge and MukF. Whereas overlapping interaction sites could result in a competition for binding between ParC, the hinge, and MukF that would affect the ATPase activity, the recent cryo-EM structure of the MukBEF complex (23), shows that in the folded conformation, the MukB hinge contacts one MukB protomer at the neck (κMukB), whereas MukF contacts the other protomer in the MukB dimer (νMukB) (Figs. 1A and 11A). This structure makes it much more likely that ParC gets sandwiched between the MukB hinge and head domains. In such a conformation, the presence of a huge ParC molecule between the hinge and head domains would inhibit their interaction, which we find important for the ATPase activity. Moreover, the crosslinking of ParC to the MukB K1314 residue present in the larynx, which opens to allow the entry of DNA into the MukB lumen, suggests that ParC could block the translocation of MukB on DNA directly as well.

### MukF Interacts with the Neck of MukB to Stimulate ATPase Activity

Our picture of the organization of the ATP hydrolysis core of the MukBEF complex has been based largely on the crystal structure of the *H. ducreyi* MukBEF complex (11). This structure showed non-hydrolyzable ATPγS sandwiched between the MukB head domains and the C-terminal winged helix domain of MukF, likely facilitating dimerization of the head domains by interacting with the cap of one of the MukB heads. Although the structure lacked the MukF N-terminal domain in complex with MukB, studies with other SMC complexes showed that the kleisin also interacted with the neck region of the SMC protein (39,40), suggesting the possibility of a similar interaction between MukB and MukF. Indeed, pulldown experiments with fragments of MukF demonstrated such an interaction between the MukF N-terminal region and the neck of MukB (41). Moreover, that study also suggested independent regulatory roles of the MukF N- and C-terminal domains in the MukBF ATPase activity. In agreement with these studies, we have found crosslinks that indicate interaction of MukF with the neck of MukB and that these interactions are vital for the stimulation of ATPase activity.

The stimulation of MukB^DNA^ ATPase activity by MukE could be because of its ability to stabilize MukF in the MukBEF complex. The mutations in MukB^DNA^ (R187E R189E), were made based on comparison to residues in the head region of the *B. subtilis* SMC protein that were required for DNA binding (27). The cryo-EM structure of Burmann et al. (23) shows R187 and R189 to be just adjacent to the DNA binding site occupied by the short duplex DNA. Given their position, these residues could interact with DNA in the absence of MukF. In the presence of MukF, these residues are likely masked by the N-terminal residues of MukF. Charge reversal substitutions at these sites could prevent binding of MukF and possibly destabilize an important interaction that decreases the efficiency of stimulation. This possibility could also explain the observations that MukF decreases the binding of MukB to DNA (42).

### Structural Bases for the Effects of Mutagenesis on the MukBEF ATPase Activity

The cryo-EM structures of Burmann et al. (23) shows the organization of the MukBEF-AcpP complex (core complex) and conformational changes upon association with ATP, MatP, and DNA (holo complex). The folded conformation of MukBEF also corresponds to crosslinking results reported previously (23) and in our present study. In the folded structure, the MukF MD, aligns with the MukB head domain, and as our crosslinking results predicted, the top of the MukF MD faces MukB neck residues.

To understand how the residues reported in our study affect the ATPase activity of the MukB complex, we mapped these residues onto the Burmann et al. (23) MukBEF core structure (Fig. 11A). On either side of the dimeric MukB structure, three residues K1232, K1246 (both on the MukB neck) and K1395 (on the MukB head) are spaced at a regular distance marking distinct regions of interaction between the MukB hinge and MukF. On one side, the MukF MD is aligned facing all three residues on νMukB, whereas the hinge domain and proximal coiled coil regions are aligned with these residues on the other side on κMukB (Fig. 11A), supporting our crosslinking results and ruling out any competition between the MukB hinge and MukF for binding to these MukB residues.

Because the conformational changes in the MukBEF structure driven by the binding of ATP and DNA are localized in the MukB head domain and coiled coil region above it, we aligned the core MukBEF and holo-MukBEF complex structures based on overall similarity followed by a forced alignment at the region of the bend in the coiled coil. As the core complex associates with ATP, MatP and two DNA molecules, the head domains are forced to close in on the bound ATP, whereas the neck regions on either side open up to favor the closed head conformation and allow extra space for DNA. The closure of the head domain seems to be enforced from either side, with the κMukB hinge pushing on the head domains from one side, whereas on the other side, MukF moves closer to the νMukB neck and inserts two MukE subunits in the MukB lumen to push the neck outward favoring an ATP-engaged conformation of the head (Supplemental Movie 1).

In the core MukBEF structure on κMukB, K1232 and K1395 side chains are present within 5 Å of those of E657 (4.3 Å) and D716 (2.3 Å) on νMukB (Figs. 11B and C), suggesting electrostatic interactions between these residues, whereas the K1246 side chain is not close to any residue on νMukB (Fig. 11D). On νMukB, however, the K1246 side chain is 5.3 Å away from the MukF D227 side chain (Fig. 11E), and these two residues could presumably close in further in a dynamic situation to stabilize the interaction between MukB and MukF. On the other hand, residues K1232 and K1395 on νMukB are far from any MukF residue (Figs. 11F and G) and thus unlikely to form such an interaction. In this structure, although νMukB K653 does face the κMukB coiled coil above the neck, we were unable to locate any interactions. However, in the holo-complex structure, νMukB K653 moves significantly closer (5.8 Å) to Glu1224 (Fig. 11H).

The mapping of these residues allowed us to classify the effect of these mutations. The observed decrease in the MukBF ATPase activity upon mutations in K1232 and K1395 likely result from a perturbed hinge and head interaction in κMukB, whereas the decrease observed with mutations at K1246 likely result from defects in interaction with MukF. Based on its position, νMukB K653 is likely interacting with the coiled coil of κMukB. Similarly, residues K283 and R286 of MukF were mapped and analyzed on these structures. In the core complex, these residues are too far away to contact any residues of MukB. Although these MukF residues move significantly toward the MukB neck in the holo-complex structure, they remain at a distance from MukB residues (Fig. 11I) However, because MukF K283 crosslinked with MukB K1232 and both residues remain in the same plane in the holo-complex structure, further inward movement of MukF in a dynamic structure would allow this contact. The interaction between MukF and MukB in this region is further substantiated by a reduced effect of MukF^2X^ on the ATPase activity of MukB K1246E. Because MukB K1246 is expected to be involved in the MukF interaction, its presence in combination with MukF^2X^ is likely to further reduce a dynamic interaction between MukF and MukB that is essential for ATPase activity. Because the decreased ATPase activities observed with other MukB mutants are likely because of decreased associations with the hinge region, they do not exhibit a significant decrease in stimulation with MukF^2X^.

### Coupling of MukB DNA Condensation to its ATPase Activity

Dependence of the SMC complexes on ATP hydrolysis for DNA organization activities and the presence of DNA and ATP binding sites on the same head domains suggests that DNA binding may affect the ATPase activity (39). Indeed, DNA does affect the ATPase activities of different SMC complexes: The *B. subtilis* SMC complex displays a DNA-dependent ATPase (43), whereas human cohesin (15) and yeast condensin (44) exhibit DNA-stimulated ATPase activities. In contrast, the effect of DNA on the MukBEF ATPase has been problematic. It has been reported that the MukB ATPase activity is DNA-independent and is, in fact, inhibited somewhat by double-stranded DNA (31), whereas Zawadzka et al. (41) showed that DNA neutralized to some extent the effect of MukE-mediated inhibition of the MukBF ATPase activity. However, in our current study, using saturating concentrations of AcpP, which is required for the MukBF ATPase activity (25), we find no effect of DNA on the ATPase activity of either the MukBF or MukBEF complexes. Nevertheless, MukB-mediated chromosomal organization is dependent on ATP hydrolysis (17).

Some of our observations thus raise the question of how presumed DNA translocation/loop extrusion by MukB is coupled to its ATPase activity. All of the variant MukB’s mutated in the head, neck, and larynx of the protein examined herein exhibited a 30%-35% decrease in ATPase activity compared to the wild type protein, yet only one of ten mutants failed to complement a Δ*mukB* strain. The MukB^DNA^ variant, which has half the ATPase activity and one-fourth the affinity for DNA as the wild type (27), also complemented a Δ*mukB* strain, yet the ATPase activity of this protein was stimulated, rather than inhibited, by MukE. Answers may lie with further analysis of the properties of the MukE^6X^ mutant, which does not inhibit MukBF ATPase activity to the same extent as the wild type MukE and does not complement a Δ*mukE* strain. Failure of MukE^6X^ to complement might arise from the inability of the MukBEF complex to translocate or localize on DNA even though the complex could hydrolyze ATP. Such a scenario, if extent, would suggest that MukE is the coupling factor between the MukBF ATPase cycle and DNA condensation.

## Experimental procedures

### Expression Plasmids

Wild-type MukB and MukB variants were either expressed in untagged form from pET11a-*mukB* (7) or with N-terminal His10-Smt3 tags from pET28b-His10-Smt3-*mukB*. ORFs for the *mukB* variants were generated by site directed mutagenesis using the Pfu Ultra II Fusion HS DNA Polymerase kit form Agilent. Primers for generating the variant ORFs are listed in Table S5. Some mutated regions of *mukB* ORFS were transferred from pET28b-Smt3-His10 plasmids to pET11a-*mukB* plasmids for complementation studies by recovering a SgrA1- and RsrII-digested plasmid from the former vector and ligating it to the latter that had also been digested with the same restriction enzymes. MukE and MukE^6X^ were expressed from pET11a-*mukE*. MukF and MukF^2X^ were expressed from pET11a-*mukF* as described (27). Primers for generating the variant ORFs are listed in Table S4. AcpP was expressed with an N-terminal Smt3-His10 tag. The *acp* ORF was amplified from pET29a-*acp*-His (Addgene) using primers listed in Table S5 and inserted into SacI- and BamH1-digested pET28b-Smt3-His10 to give pET28b-Smt3-His10-*acp*.

### Proteins

Proteins were expressed in *E. coli* BL21(DE3) as described (7,27). Untagged MukB wild type and variants were purified as described (7). Tagged MukB proteins were purified as follows (all procedures were at 4 °C): Cells were lysed with lysozyme and MukB proteins were precipitated by the addition of (NH_4_)_2_S0_4_ to the cleared S100 lysates to 32% saturation. Protein pellets were resuspended in Buffer T (50 mM potassium phosphate (pH 8.0), 300 mM NaCl, 0.5% Triton X-100, and 20% glycerol) and dialyzed overnight in the same buffer. The dialyzed fraction was mixed for 3 h with Talon Metal Affinity Resin (Clontech) that had been equilibrated with Buffer T. The slurry was packed into a column and washed with 20 column volumes of Buffer T + 25 mM imidazole-HCl (pH 8.0). MukB proteins were eluted with Buffer T + 0.5 M imidazole-HCl and incubated overnight with Ulp1 at a 1:50 ratio (w/w, Ulp1:MukB) to cleave off the tag. Products were precipitated with (NH_4_)_2_S0_4_ at 50% saturation, resuspended in Buffer A (50 mM Tris-HCl (pH 7.5), 5 mM DTT, 1 mM EDTA, 0.5 M NaCl, and 20% glycerol), and gel filtered through a Superose 6 10/300 GL column (GE Healthcare) equilibrated with the same buffer. Peak fractions (located by SDS-PAGE) were pooled, dialyzed overnight against storage buffer (50 mM Tris-HCl (pH 7.5), 5 mM DTT, 1 mM EDTA, 0.2 M NaCl, and 40% glycerol), aliquoted, frozen in liquid N_2_, and stored at -80 °C. All final fractions were free of detectable nuclease activity. Wild-type MukB purified either without or via a tag had identical ATPase activity (Fig. S2). MukB wild type and variants K653A, K653E, K1246A, and K1246E were purified without a tag. MukB wild type and variants K1232A, K1232E, K1314A, K1314E, K1395A, and K1395E were purified via a tag. Fig. S3 shows the different MukB preparations and their saturation with AcpP. Saturation by AcpP could vary considerably, as illustrated by the gel of the two wild-type MukB preparations shown in Fig. 1A.

MukB hinge fragment. The MukB hinge ORF (residues 645-804) was expressed from pRSFDuet with an N-terminal His-Smt3 tag in BL21(DE3). Three liters of cells were grown at 37 ° C in LB medium containing 50 μg/ml kanamycin to O.D._600_ = 0.5, cooled to 16 °C, and protein expression induced by the addition of IPTG to 0.4 mM. Cells were harvested after 16 h, resuspended in Buffer A (50 mM Tris HCl (pH 8.0), 500 mM NaCl, 2 mM PMSF, 5 % (v/v) glycerol) containing 10 mM imidazole, lysed by sonication, and the cleared lysate bound to Ni-NTA resin (Invitrogen) overnight at 4 °C. The resin was washed with buffer A containing 50 mM imidazole and protein was eluted using 300 mM imidazole in buffer A. Peak fractions (located by SDS-PAGE) were pooled, digested with Ulp1 overnight while being dialyzed against buffer A containing 10 mM imidazole, and passed through Ni-NTA resin again to remove the cleaved tag and Ulp1. Protein in the flow through was precipitated by the addition of (NH_4_)SO_4_ to 55 % saturation, dissolved in buffer A, and gel filtered through a 135 ml Superdex 200 column developed with Buffer A. Peak fractions were pooled, dialyzed against storage buffer (50 mM Tris HCl pH 7.5, 150 mM NaCl, 2 mM DTT, 1 mM EDTA and 40 % glycerol) overnight, aliquoted, frozen in liquid N_2_, and stored at -80 °C. SDS-PAGE analysis of MukB hinge fragment is shown in Fig. S4.

MukE and MukE^6X^ were purified as described (27) with the exception that the order of the gel filtration and hydroxyapatite chromatography steps were reversed. SDS-PAGE analysis of MukE^6X^ is shown in Fig. S4.

MukF and MukF^2X^ were purified as described (27) except that fraction 3 was precipitated with (NH_4_)_2_S0_4_ at 35% saturation rather than 60%. SDS-PAGE analysis of MukF^2X^ is shown in Fig. S4.

ParC was purified as described (45).

AcpP. Eight liters of BL21(DE3)(pET28b-Smt3-His10-*acp*) in LB medium + 50 μg kanamycin were grown at 37 °C to an O.D._600_ of 0.8, IPTG was added to 1 mM, and growth continued overnight at 16 °C. Cells were harvested, resuspended in 50 mM Tris-HCl (pH 8.0 at 4 °C), 0.5 M NaCl, 10 mM imidazole-HCl (pH 8.0), 10% sucrose, 5% glycerol, and 1X Bacterial Protease Arrest (G-biosciences) to 150 O.D._600_/ml, and lysed by three passages through a French Pressure Cell at 20,000 psi. A cleared S100 lysate was mixed for 3 h with Ni-NTA resin (Invitrogen) equilibrated with Buffer B (50 mM Tris-HCl (pH 8.0 at 4 °C), 0.5 M NaCl, and 5% glycerol) containing 10 mM imidazole-HCl (pH 8.0). The slurry was then poured into a disposable column, washed with 20 column volumes of Buffer B containing 10 mM imidazole-HCl (pH 8.0), followed by five column volumes of Buffer B containing 20 mM imidazole-HCl (pH 8.0). AcpP was eluted with Buffer B containing 100 mM imidazole-HCl (pH 8.0), incubated overnight with Ulp1 at a 1:70 ratio (w/w, Ulp1:AcpP) to cleave off the tag, and dialyzed overnight against Buffer B containing 10 mM imidazole-HCl (pH 8.0) and 0.05% NP-40. The dialyzed fraction was mixed for 2 h with Ni-NTA resin (Invitrogen) equilibrated with the same buffer, poured into a disposable column, and the flow through was then gel filtered through a Superdex 200 (130 ml) column equilibrated in 50 mM Tris-HCl (pH 8.0 at 4 °C), 1 mM EDTA, 5 mM DTT, 300 mM NaCl, 0.05% NP-40, and 10% glycerol. Peak fractions (located by SDS-PAGE) were pooled, dialyzed against storage buffer (50 mM Tris-HCl (pH 8.0 at 4 °C), 1 mM EDTA, 5 mM DTT, 150 mM NaCl, 0.05% NP-40, and 40% glycerol), aliquoted, frozen in liquid N_2_, and stored at -80 °C.

### Protein-Protein Crosslinking Analyses

Proteins were dialyzed extensively against 50 mM HEPES-NaOH (pH 8.0), 150 mM KCl, and 20% glycerol to remove DTT and EDTA. For MukB alone, MukB (125 nM as dimer) was incubated with 40 μM disuccinimdyl glutarate in conjugation buffer (50 mM HEPES-NaOH (pH 8.0), 0.5 mM Mg(OAC)_2_, and 20 mM KCl) at 37 °C for 20 min, the reaction was quenched by the addition of Tris-HCl (pH 8.8) to 75 mM followed by incubation at room temperature for 5 min, and the products precipitated by the addition of TCA to 5%. Proteins were recovered by centrifugation, dissolved in SDS-PAGE loading dye, and electrophoresed through a NuPAGE 3%-8% Tris-Acetate gel. The gel was stained with Simply Blue SafeStain (Invitrogen), species with mobilities slower than monomeric MukB were excised, and the samples submitted to the MSK Mass Spectrometry Core. Samples were processed as described (46). For MukB and ParC, 0.9 μM MukB and 1.8 μM ParC (as dimers) were treated with 90 μM DSG. For MukB and MukF, 0.75 μM MukB and 2.5 μM MukF (as dimers) were treated with 200 μM DSG.

The LC-MS/MS .raw files were processed using Mascot (version 2.6.1.100) and searched for protein identification against the SwissProt protein database for *E. coli*. Carbamidomethylation of C was set as a fixed modification and the following variable modifications allowed: oxidation (M), N-terminal protein acetylation, deamidation (N and Q), ubiquitination (K), and phosphorylation (S, T and Y). Search parameters specified an MS tolerance of 10 ppm, an MS/MS tolerance at 0.08 Da and full trypsin digestion, allowing for up to two missed cleavages. False discovery rate was restricted to 1% in both protein and peptide level. Protein coverage and peptide count were obtained using Scaffold (version 4.8.4).

### Mass Spectrometry Analysis of the MukB Hinge Fragment

MukB hinge fragment (26 μg) was denatured in 8 M urea (200 ml), treated with 5 mM TCEP at 56 °C for 20 min, and alkylated with 10 mM 2-iodoacetamide at room temperature for 30 min in the dark. The reaction was quenched by the addition of DTT to 5 mM followed by incubation at room temperature for 15 min. Tris-HCl (pH 8) was then added to 4 mM and the sample divided into two equal portions. One portion was treated with 0.5 μg of trypsin at 37 °C and the other with 0.5 μg of chymotrypsin at room temperature overnight. Samples were then desalted and analyzed by mass spectrometry.

### ATPase Assays

The ENZCheck Phosphate Assay Kit (Life Technologies) was used. Reaction mixtures (150 μl) containing the supplied standard reaction buffer supplemented with 2 mM ATP, 20 mM KCl, and 1 mM Mg(OAC)_2_ and the indicated concentrations of proteins were assembled on ice, aliquots (140 μl) were transferred to a chilled 96 well plate (Corning 9017 with lid), the plate shaken for 2 s at 180 rpm, and the reaction analyzed immediately in a BioTek Epoch 2 Microplate Reader preheated to 37 °C. Optical density at 330 and 360 nm was recorded at 2 min intervals for 30 min. A sample that did not contain proteins was used as the blank in all experiments.

### Complementation Assays

pET11a plasmids carrying wild-type and mutated *mukB, mukE*, and *mukF* alleles were transformed into BW30270*ΔmukB::kan* (10), AZ5381 *mukF::kan* trpC9941 (6), and AZ5450 *mukE::kan* trpC9941 (6). Post-transformation recovery was at 25 °C. Single colonies from these transformants were grown overnight in LB medium containing 100 μg/ml ampicillin at 25 °C, sub-cultured the next day to 0.05 O.D._600_ in 50 ml medium, grown to 0.6 O.D._600_, diluted serially in phosphate buffered saline, and plated on LB agar plates containing ampicillin that were then grown at 25 °C (permissive temperature) and 37 °C (non-permissive temperature). Protein expression from the plasmids was confirmed by harvesting transformed cells grown at 25 °C to mid-log phase, resuspending in lysis buffer (50 mM Tris-HCl (pH 7.5), 150 mM NaCl, 1 mM EDTA, 2 mM DTT, and 5 % glycerol), lysing by sonication, and Western blotting varying equivalent amounts of cell lysate resolved by SDS-PAGE. Blots were probed with affinity-purified MukB, MukE, and MukF polyclonal antibodies as indicated. Blots were developed using HRP-conjugated secondary antibodies and ECL reagents (Cytiva Life Sciences).

### Data availability

All raw data and images are held by the authors and are available upon request.

## Supporting information

This article contains supporting information.

## Footnotes

## 1, the abbreviations used are

SMC: structural maintenance of chromosomes
Topo IV: topoisomerase IV
MD: middle domain
apo-MukB (MukB^apo^): referring to the MukB dimer alone with no associated AcpP
holo-MukB (MukB^holo^): referring to the stoichiometric complex of the MukB dimer with AcpP
core MukBEF complex: referring to the complex of MukB-AcpP, MukE, and MukF
holo-MukBEF complex: referring to the complex of MukB-AcpP, MukE, MukF, MatP, DNA, and ATP

