## Supplemental Information for "Inter- and intra-Subunit interactions driving the MukBEF ATPase"

Running title: MukB ATPase

Keywords: ATPase, DNA binding protein, DNA structure, genome structure, nucleic acid enzymology

Contents

Tables S1-S5, note that tables S1-S3 have been submitted independently as .xls files

Figures S1-S4

Table S4. Mass Spectrometry Analysis of MukB Hinge Preparation

**Probability Legend**

- over 95%
- 80% to 94%
- 50% to 79%
- 20% to 49%
- 0% to 19%

| # | BioView: 35 Proteins in 35 Clusters | Accession Number | Molecular Weight | Protein Grouping Ambiguity | 1 S2_A1_Trypsin | 2 S2_A1_Chymo |
|---|---|---|---|---|---|---|
| 1 | Chromosome partition protein MukB OS=Escherichia coli O1:K1 / APEC GN=mukB PE=3 SV=1 | sp\|A1A9K1\|MUKB_ECOK1 (+20) | 170 kDa | | 1632 | 333 |
| 2 | ATP-dependent RNA helicase RhlE OS=Escherichia coli (strain K12) GN=rhlE PE=1 SV=3 | sp\|P25888\|RHLE_ECOLI | 50 kDa | | 5 | 2 |
| 3 | ATP-dependent protease ATPase subunit HslU OS=Escherichia coli O1:K1 / APEC GN=hslU PE=3 SV=1 | sp\|A1AIA6\|HSLU_ECOK1 (+20) | 50 kDa | | 2 | |
| 4 | Biotin carboxylase OS=Escherichia coli (strain K12) GN=accC PE=1 SV=2 | sp\|P24182\|ACCC_ECOLI (+1) | 49 kDa | | 27 | |
| 5 | Transcription termination factor Rho OS=Escherichia coli (strain K12) GN=rho PE=1 SV=1 | sp\|P0AG30\|RHO_ECOLI (+2) | 47 kDa | | 6 | |
| 6 | Phosphopentomutase OS=Escherichia coli O139:H28 (strain E24377A / ETEC) GN=deoB PE=3 SV=1 | sp\|A7ZVS6\|DEOB_ECO24 (+17) | 44 kDa | | 15 | 3 |
| 7 | Elongation factor Tu 1 OS=Escherichia coli O1:K1 / APEC GN=tuf1 PE=3 SV=2 | sp\|A1AGM6\|EFTU1_ECOK1 (+15) | 43 kDa | | 9 | |
| 8 | HTH-type transcriptional regulator GalR OS=Escherichia coli (strain K12) GN=galR PE=1 SV=1 | sp\|P03024\|GALR_ECOLI | 37 kDa | | 3 | 2 |
| 9 | Aminoglycoside 3'-phosphotransferase OS=Escherichia coli GN=aphA1 PE=3 SV=2 | sp\|P00551\|KKA1_ECOLX | 31 kDa | | 13 | 2 |
| 10 | 50S ribosomal protein L2 OS=Escherichia coli O1:K1 / APEC GN=rplB PE=3 SV=1 | sp\|A1AGK5\|RL2_ECOK1 (+20) | 30 kDa | | 26 | 1 |
| 11 | HTH-type transcriptional repressor BluR OS=Escherichia coli (strain K12) GN=bluR PE=1 SV=1 | sp\|P75989\|BLUR_ECOLI | 28 kDa | | 3 | |
| 12 | 30S ribosomal protein S3 OS=Escherichia coli O1:K1 / APEC GN=rpsC PE=3 SV=1 | sp\|A1AGK2\|RS3_ECOK1 (+19) | 26 kDa | | 48 | 3 |
| 13 | tRNA (guanosine(18)-2'-O)-methyltransferase OS=Escherichia coli (strain K12) GN=trmH PE=1 SV=1 | sp\|P0AGJ2\|TRMH_ECOLI (+1) | 25 kDa | | 2 | |
| 14 | Protein InaA OS=Escherichia coli (strain K12) GN=inaA PE=1 SV=3 | sp\|P27294\|INAA_ECOLI | 25 kDa | | 7 | |
| 15 | 50S ribosomal protein L1 OS=Escherichia coli O139:H28 (strain E24377A / ETEC) GN=rplA PE=3 SV=1 | sp\|A7ZUJ7\|RL1_ECO24 (+19) | 25 kDa | | 2 | |
| 16 | 30S ribosomal protein S4 OS=Escherichia coli O1:K1 / APEC GN=rpsD PE=3 SV=1 | sp\|A1AGI7\|RS4_ECOK1 (+20) | 23 kDa | | 21 | 2 |
| 17 | 50S ribosomal protein L3 OS=Escherichia coli O1:K1 / APEC GN=rplC PE=3 SV=1 | sp\|A1AGK7\|RL3_ECOK1 (+20) | 22 kDa | | 3 | |
| 18 | Alkyl hydroperoxide reductase subunit C OS=Escherichia coli (strain K12) GN=ahpC PE=1 SV=2 | sp\|P0AE08\|AHPC_ECOLI (+2) | 21 kDa | | 5 | 3 |
| 19 | 50S ribosomal protein L5 OS=Escherichia coli O1:K1 / APEC GN=rplE PE=3 SV=1 | sp\|A1AGJ7\|RL5_ECOK1 (+20) | 20 kDa | | 2 | 1 |
| 20 | 30S ribosomal protein S5 OS=Escherichia coli O1:K1 / APEC GN=rpsE PE=3 SV=1 | sp\|A1AGJ2\|RS5_ECOK1 (+8) | 18 kDa | | 17 | 6 |
| 21 | Ferric uptake regulation protein OS=Escherichia coli (strain K12) GN=fur PE=1 SV=1 | sp\|P0A9A9\|FUR_ECOLI (+2) | 17 kDa | | 5 | 2 |
| 22 | Biotin carboxyl carrier protein of acetyl-CoA carboxylase OS=Escherichia coli (strain K12) GN=accB PE=1 SV=1 | sp\|P0ABD8\|BCCP_ECOLI (+2) | 17 kDa | | 4 | 2 |
| 23 | 50S ribosomal protein L9 OS=Escherichia coli O1:K1 / APEC GN=rplI PE=3 SV=1 | sp\|A1AJA7\|RL9_ECOK1 (+20) | 16 kDa | | 14 | |
| 24 | 50S ribosomal protein L16 OS=Escherichia coli O1:K1 / APEC GN=rplP PE=3 SV=1 | sp\|A1AGK1\|RL16_ECOK1 (+20) | 15 kDa | | 5 | 7 |
| 25 | 50S ribosomal protein L15 OS=Escherichia coli O1:K1 / APEC GN=rplO PE=3 SV=1 | sp\|A1AGJ1\|RL15_ECOK1 (+20) | 15 kDa | | 6 | 1 |
| 26 | 50S ribosomal protein L11 OS=Escherichia coli O139:H28 (strain E24377A / ETEC) GN=rplK PE=3 SV=1 | sp\|A7ZUJ6\|RL11_ECO24 (+19) | 15 kDa | | 3 | |
| 27 | 30S ribosomal protein S9 OS=Escherichia coli O1:K1 / APEC GN=rpsI PE=3 SV=1 | sp\|A1AGC3\|RS9_ECOK1 (+20) | 15 kDa | | 11 | |
| 28 | 50S ribosomal protein L17 OS=Escherichia coli O1:K1 / APEC GN=rplQ PE=3 SV=1 | sp\|A1AGI5\|RL17_ECOK1 (+19) | 14 kDa | | 10 | 4 |
| 29 | 30S ribosomal protein S11 OS=Escherichia coli O1:K1 / APEC GN=rpsK PE=3 SV=1 | sp\|A1AGI8\|RS11_ECOK1 (+20) | 14 kDa | | 8 | 2 |
| 30 | 30S ribosomal protein S12 OS=Escherichia coli O1:K1 / APEC GN=rpsL PE=3 SV=1 | sp\|A1AGM9\|RS12_ECOK1 (+18) | 14 kDa | | 8 | 1 |
| 31 | 50S ribosomal protein L18 OS=Escherichia coli O1:K1 / APEC GN=rplR PE=3 SV=1 | sp\|A1AGJ3\|RL18_ECOK1 (+20) | 13 kDa | | 7 | 2 |
| 32 | 30S ribosomal protein S14 OS=Escherichia coli O1:K1 / APEC GN=rpsN PE=3 SV=1 | sp\|A1AGJ6\|RS14_ECOK1 (+15) | 12 kDa | | 2 | 2 |
| 33 | 50S ribosomal protein L21 OS=Escherichia coli O139:H28 (strain E24377A / ETEC) GN=rplU PE=3 SV=1 | sp\|A7ZS83\|RL21_ECO24 (+18) | 12 kDa | | 3 | 1 |
| 34 | 50S ribosomal protein L24 OS=Escherichia coli O139:H28 (strain E24377A / ETEC) GN=rplX PE=3 SV=1 | sp\|A7ZSJ8\|RL24_ECO24 (+13) | 11 kDa | | | 2 |
| 35 | 30S ribosomal protein S15 OS=Escherichia coli O1:K1 / APEC GN=rpsO PE=3 SV=1 | sp\|A1AG70\|RS15_ECOK1 (+18) | 10 kDa | | 2 | |

Thirteen micrograms of purified MukB hinge fragment preparation were digested with either trypsin or chymotrypsin and analyzed by mass spectrometry as described under Experimental Procedures. Shown is the “Total Spectrum Count” analysis from Scaffold (v.4.8.4).

Table S5. Oligonucleotides used in this study

Primers for generating a BamHI-SacI insert fragment of MukB

| | |
|---|---|
| MukB-BamHI | GAA GGA GAT **GGA TCC** ATG ATT GAA CGC GGT AAA TTT CGC TCA CTG |
| MukB-SacI | TTT TTT TTT T**GA GCT C**TT AAC TCG CCT GAG AAG GCG CTT CGT CAG |

Site Directed Mutagenesis Primers for MukB:

| | |
|---|---|
| K653A 5’ | GAACGCGATGAAGTGGGCGCGCGCGCAAACGCCGTCGATGAAGAGATCG |
| K653A 3’ | CGATCTCTTCATCGACGGCGTTTGCGCGCGCGCCCACTTCATCGCGTTC |
| K653E 5’ | GAACGCGATGAAGTGGGCGCGCGCGAAAACGCCGTCGATGAAGAGATCG |
| K653E 3’ | CGATCTCTTCATCGACGGCGTTTTCGCGCGCGCCCACTTCATCGCGTTC |
| K1232A 5’ | CTCCCGTGAACAGGCACTGGCGATCAGTTC |
| K1232A 3’ | GAACTGATCGCCAGTGCCTGTTCACGGGAG |
| K1232E 5’ | CTCCCGTGA ACAGGAACT GGCGATCAGTTC |
| K1232E 3’ | GAACTGATCGCCAGTTCCTGTTCACGGGAG |
| K1246A 5’ | CGCAGCGTGGCGAACATCATTCGCGCAACCATTCAGCGCGAGCAGAACCG |
| K1246A 3’ | CGGTTCTGCTCGCGCTGAATGGTTGCGCGAATGATGTTCGCCACGCTGCG |
| K1246E 5’ | CGCAGCGTGGCGAACATCATTCGCGAAACCATTCAGCGCGAGCAGAACCG |
| K1246E 3’ | CGGTTCTGCTCGCGCTGAATGGTTTCGCGAATGATGTTCGCCACGCTGCG |
| K1314A 5’ | GGAAGCGCTGGCGGCACTGTATC ACGTC |
| K1314A 3’ | GACGTTGATACAGTGCCGCCAGCGCTTCC |
| K1314E 5’ | GGAAGCGCTGGCGGAACTGTATCAACGTC |
| K1314E 3’ | GACGTTGATACAGTTCCGCCAGCGCTTCC |
| K1395A 5’ | CCGCCTGCGCGGTGCAGATATCTCTCCTTG |
| K1395A 3’ | CAAGGAGAGATATCTGCACCGCGCAGGCGG |
| K1395E 5’ | CCGCCTGCGCGGTGAAGATATCTCTCCTTG |
| K1395E 3’ | CAAGGAGAGATATCTTCACCGCGCAGGCGG |

AcpP ORF Primers:

| | |
|---|---|
| AcpP-SacI | 5’-TTT TTT TTT **GAG CTC** TTA CGC CTG GTG GCC GTT GAT GTA ATC |
| AcpP-BamHI | 5’-TTT TTT TTT **GGA TCC** AGC ACT ATC GAA GAA CGC GTT AAG |

50mer ds DNA:

5' - ATC TCG GGC TAT TCT TTT GAT TTA TAA GGG ATT TTG CCG ATT TCG GAA CC

3' - GG TTC CGA AAT CGG CAA AAT CCC TTA TAA ATC AAA AGA ATA GCC CGA GAT

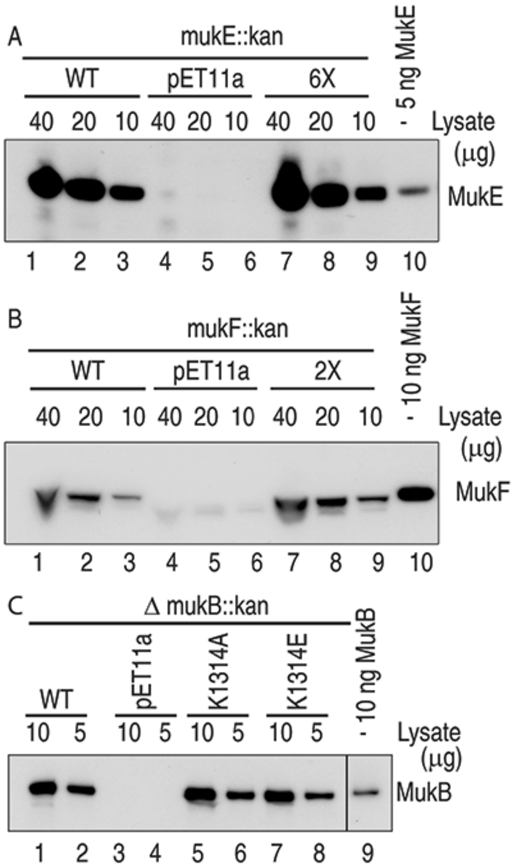


Fig. S1. Western blot expression analyses of various *mukB*, *mukF*, and *mukE* mutant alleles. A, *mukE* and *mukE*$^{6X}$ expressed from pET11a in AZ5450 at 25 ºC and analyzed as described under Experimental Procedures. 6X, *mukE*$^{6X}$. B, *mukF* and *mukF*$^{2X}$ expressed from pET11a in AZ5381 at 25 ºC and analyzed as described under Experimental Procedures. 2X, *mukF*$^{2X}$. C, *mukB*, *mukBK1314A*, and *mukBK1314E* expressed from pET11a in BW30270*ΔmukB::kan* at 25 ºC and analyzed as described under Experimental Procedures.

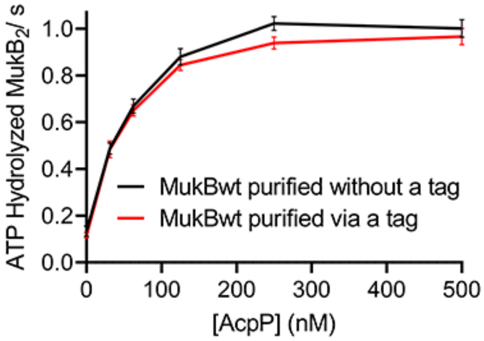


Fig. S2. MukB purified without a tag or via a His-Smt3 tag that was then cut off have identical ATPase activity. The indicated proteins were assayed in standard ATPase reactions containing either no or 32 nM, 63 nM, 125 nM, 250 nM, and 500 nM AcpP. Maximal ATPase rates are plotted as a function of [AcpP].

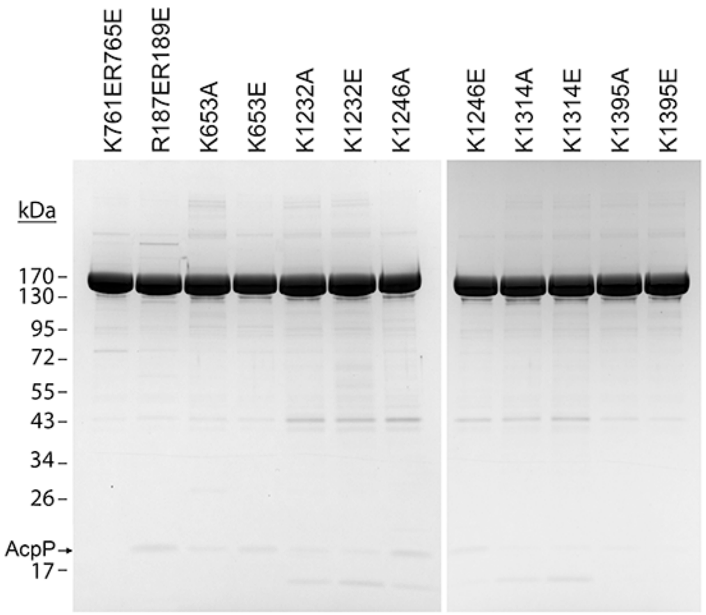


Fig. S3. SDS-PAGE of variant MukB proteins preparations used in this study.
Seven and one half micrograms of the indicated preparations were electrophoresed through a Novex 4%-20% gradient gel. The gel was stained with SimplyBlue.

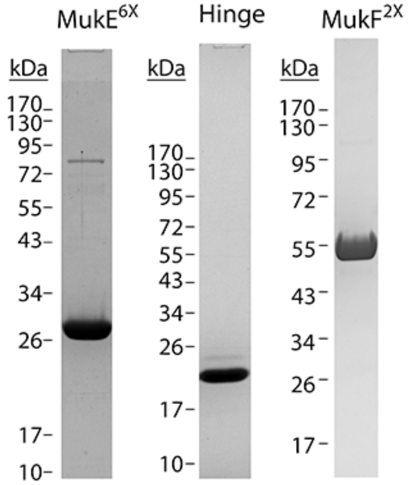


Fig. S4. SDS-PAGE of MukE$^{6X}$, MukB hinge fragment, and MukF$^{2X}$ preparations used in this study. MukE$^{6X}$: 5 $\mu$g on a Novex 12% gel. Hinge fragment: 5 $\mu$g on a Novex 4%-20% gel. MukF$^{2X}$: 5 $\mu$g on an 8% gel.
